# Tumor stiffness creates a ferroptotic Achilles’ heel through YAP–SKP2 signaling

**DOI:** 10.64898/2026.10.01.755825

**Authors:** Emma Villares, Nico Taber, Shao-Chin Wu, Connor Amelung, Jianli Wu, Chao-Chieh Lin, Jen-Tsan Ashley Chi, Sharon Gerecht

## Abstract

Matrix stiffness is a hallmark of aggressive solid tumors and strongly associated with therapeutic resistance and poor prognosis. This mechanical barrier represents a major unmet challenge in oncology, as current strategies rarely address how matrix mechanics themselves drive treatment resistance. Here, we identified ferroptosis, an iron-dependent form of cell death, as a potential long-sought vulnerability that can be exploited to eliminate mechanically resistant tumors. Using genetic and chemical approaches, we investigated how matrix stiffness sensitizes cancer cells to ferroptosis and whether this vulnerability can be therapeutically targeted. We found that cancer cells cultured on stiff matrices were more sensitive to ferroptosis than those on soft matrices, as evidenced by reduced cell viability and elevated lipid peroxidation. Furthermore, we found that matrix stiffness induced YAP activation, which upregulated its target gene *SKP2*, a key mediator required for the heightened ferroptosis sensitivity observed under stiff conditions. Finally, using an injectable hydrogel system, we demonstrated that stiffened tumors were resistant to chemotherapy but remained highly susceptible to ferroptosis induction, resulting in markedly reduced tumor volume and weight. Our findings established ferroptosis as a stiffness-dependent vulnerability mediated by a YAP-SKP2 signaling axis, revealing a mechanotransductive pathway that may be leveraged to target treatment-refractory tumors.

## Introduction

The extracellular matrix (ECM) is a dynamic network of proteins surrounding cells that has emerged as a key regulator of the tumor microenvironment, influencing cancer progression and therapeutic response (1–5). In solid tumors, the ECM is stiffer and has a distinctive composition compared to normal tissue due to dysregulated matrix remodeling (6). This results from an imbalance between ECM synthesis and degradation, driven by aberrant expression of matrix-remodeling enzymes (7, 8). Abnormal ECM in cancer affects oncogenic signaling and gene expression, promotes cancer progression and metastasis, and contributes to therapeutic resistance (9–12). Increasing evidence indicates that matrix stiffening not only promotes tumor growth and invasion but also protects cancer cells from conventional cell death pathways such as apoptosis and autophagy. These mechanical cues create a microenvironment that shields tumor cells from chemotherapeutic and targeted agents, leading to treatment-refractory phenotypes and poor patient prognosis (13, 14). Given the critical role of ECM stiffness in fostering chemoresistance and protecting tumor cells from diverse conventional treatments, there is an urgent and unmet need for therapeutic strategies that can selectively target and eliminate these mechanically adapted, treatment-refractory tumor cells. A promising direction has emerged through the recent discovery of ferroptosis, an iron-dependent form of regulated cell death driven by lipid peroxidation, distinct from apoptosis, autophagy, or necrosis (15). Ferroptosis plays a critical role in various disease contexts, including ischemia-reperfusion injury, neurodegenerative diseases, and, particularly, in cancer (16–19). This process is initiated when labile iron (Fe^2+^) catalyzes the formation of reactive oxygen species (ROS), specifically hydroxyl radicals, via the Fenton reaction, leading to lipid membrane peroxidation (20). Normally, glutathione peroxidase 4 (GPX4) neutralizes lipid peroxides by converting them to non-toxic lipid alcohols using glutathione (GSH) as a cofactor (20). However, when GPX4 is inhibited or GSH is depleted, lipid peroxides accumulate, leading to cellular damage and ultimately cell death (20, 21). Ferroptosis can be triggered by various ferroptosis-inducing compounds (FINs), which can be categorized into different classes based on their targets and mechanisms (22). The class I FINs target xCT transporters, including erastin and sulfasalazine (SAS), to block cystine uptake. Class II FINs directly inhibit the enzymatic activity of GPX4, preventing the reduction of lipid hydroperoxides. Class III FINs, such as FIN56, promote GPX4 protein degradation and coenzyme Q₁₀ (CoQ₁₀)(23). Class IV FINs, such as FINO₂, oxidize iron and promote lipid peroxidation independently of GPX4 inhibition. Together, these compounds define mechanistically distinct routes to trigger ferroptotic cell death, as shown in our recent studies on the regulation of ferroptosis by Coenzyme A(24) and NINJ1(25) that specifically affect class I FIN-induced cell death.

As recently shown, ferroptosis sensitivity is regulated by cellular contacts and density via the hippo pathway effector, YAP (Yes-associated protein) transcription and nuclear localization (18, 26). YAP is a transcriptional co-activator that is one of the primary mediators of ECM stiffness-induced signaling (27, 28). When cells experience a soft ECM, it sequesters YAP in the cytoplasm, preventing its interaction with TEAD transcription factors and thereby suppressing TEAD-dependent gene expression. However, when cells sense an increase in ECM stiffness, YAP translocates to the nucleus to cooperate with TEAD and activate transcriptional programs that promote cell proliferation, survival, and metabolic adaptation (27). This suggests a potential mechanistic link between ferroptosis sensitivity and ECM stiffness via YAP signaling. More importantly, targeting this pathway could provide a therapeutic strategy to induce ferroptosis in stiff, chemoresistant tumors, offering an avenue for overcoming treatment resistance.

Here, we demonstrate that matrix stiffness regulates ferroptosis in cancer. We first show that cancer cells cultured on stiff hydrogels exhibit significantly decreased viability and increased lipid peroxidation compared to those cultured on soft hydrogels across four different solid tumor types and in response to three ferroptosis-inducing agents with distinct mechanisms. Further, we show that inhibition of YAP, using both small-molecule inhibitors and genetic approaches, rescues ferroptosis sensitivity in cells cultured on stiff substrates. We demonstrate that stiffness-dependent differences in cell-cycle progression contribute to the observed variation in ferroptotic sensitivity. Therefore, we identified *SKP2*, a YAP target gene and cell cycle driver, as a key player in stiffness-mediated sensitivity. Knockdown of *SKP2* rescued cells cultured on stiff substrates from ferroptosis. Finally, we employed a xenograft model and demonstrated that we can exploit ferroptosis to target stiffened tumors in vivo. Overall, the presented work reveals that matrix mechanics regulate ferroptosis via the YAP-SKP2 pathway, providing an avenue for treating stiff, chemoresistant tumors.

## Results

### Matrix stiffness regulates ferroptosis in breast, ovarian, renal, and fibrosarcoma cancers

Because matrix stiffening protects tumor cells from conventional cell death pathways such as apoptosis, we reasoned that it might similarly regulate ferroptosis sensitivity. To test this possibility, we utilized stiff (25 kPa) and soft (0.5 kPa) polyacrylamide (PA) hydrogels. We selected 0.5 kPa for our soft hydrogel to reflect the stiffness of normal tissue and very soft malignancies (29). We selected 25 kPa for the stiff hydrogel, reflecting the upper range of stiffness reported for multiple tumor types—breast, fibrosarcoma, and ovarian cancer—and its link to therapeutic resistance.(6, 30) We found that cancer cells seeded onto stiff PA hydrogels showed greater sensitivity to ferroptosis than those seeded onto soft PA hydrogels. Cell viability assays across multiple cancer cell lines of both mesenchymal and epithelial origin, including MDA-MB-231 (breast cancer), RCC4 (renal cell carcinoma), CAOV3 (ovarian cancer), and HT1080 (fibrosarcoma), were performed after treatment with erastin, a class I ferroptosis-inducer that targets the upstream xCT glutamate-cysteine antiporter. In each case, erastin-treated cells on stiff hydrogels showed significantly reduced viability compared with those on soft hydrogels, suggesting that the impact of matrix stiffness on ferroptosis sensitivity is a consistent phenomenon across various solid tumor types (Figure 1A).

**Figure 1:**
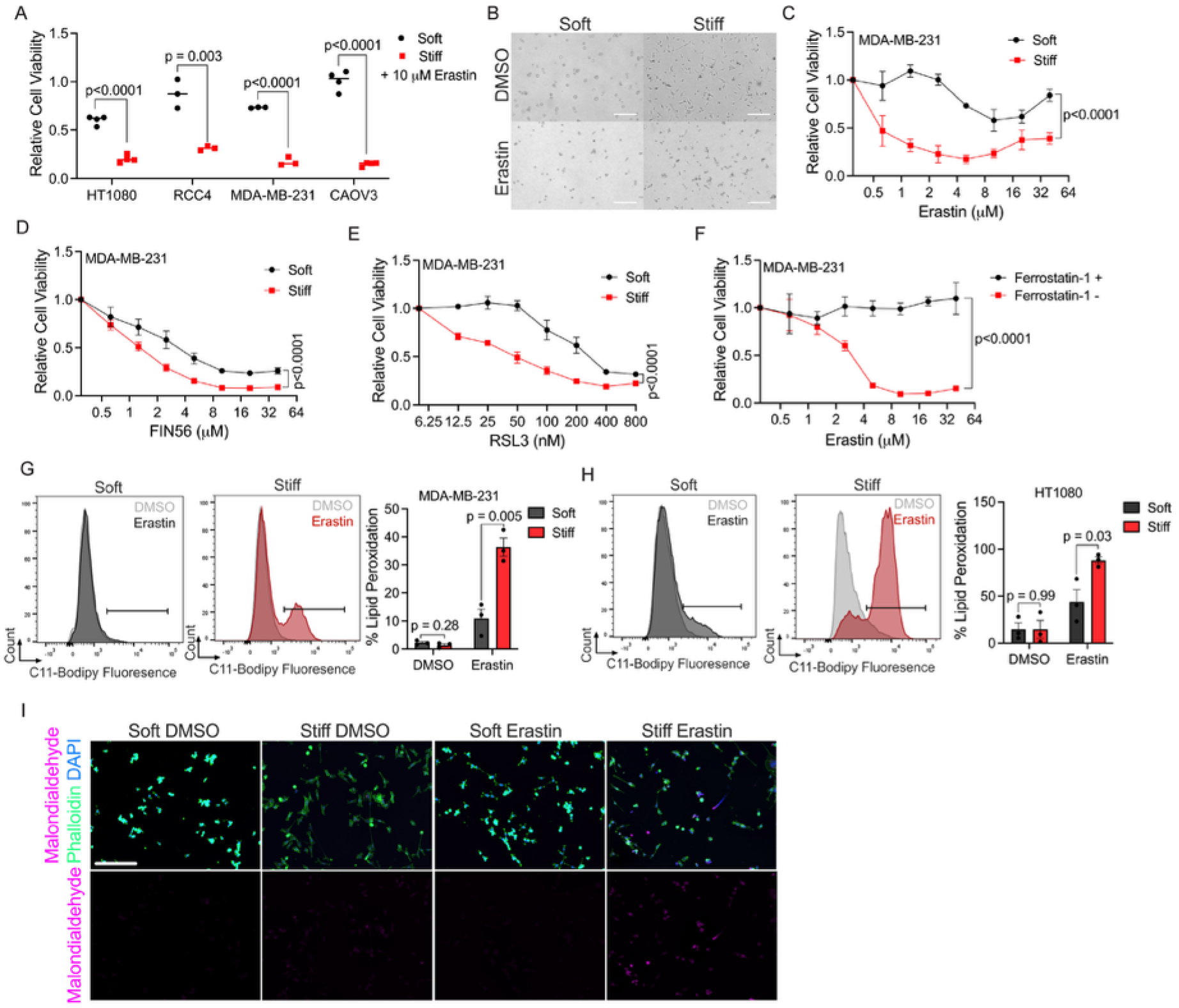
Ferroptosis is regulated by matrix stiffness in multiple cancer types. (**A**) Relative cell viability of multiple cancer cell lines cultured on soft or stiff hydrogels following 16hr treatment with 10 μM erastin. (HT1080 N=4, RCC4 N=3, MDA-MB-231 N=3, CAOV3 N=4) (**B**) Brightfield images of MDA-MB-231 cells on soft and stiff hydrogels treated with 10 μM erastin or DMSO for 16 h. Scale bar=100μm (**C**) Relative cell viability of MDA-MB-231 breast cancer cells on soft or stiff hydrogels treated with erastin (N=4), (**D**) FIN56, or (**E**) RSL3 for 16 hr. (N=3) (**F**) MDA-MB-231 cells on stiff hydrogels treated with erastin with or without 6 μM ferrostatin-1 (ferroptosis inhibitor) for 16 hr. (N=3) (**G**) Representative flow cytometry histograms of lipid peroxidation measured by C11 Bodipy fluorescence of MDA-MB-231 cells treated with either 10 μM erastin or DMSO control on soft or stiff hydrogels for 6 hr (left) and the quantification (right; N=3). (**H**) Representative flow cytometry histograms of lipid peroxidation measured by C11 Bodipy fluorescence of HT1080 cells treated with either 10 μM erastin or DMSO control on soft or stiff hydrogels for 6 hr (left) and the quantification (right; N=3). (**I**) Representative immunofluorescence images of MDA-MB-231 cells stained with malondialdehyde, phalloidin, and DAPI treated with 10 μM erastin or DMSO for 12 hr of N=3 independent experiments. Scale bar=100μm. All graphs represent means ± SEM except 1A where lines in graphs represent mean. One-way ANOVA was used for A, G, H. Two-way repeated-measures ANOVA was used for C-F.

These results were further validated by morphological observations. MDA-MB-231 breast cancer cells cultured on stiff substrates exhibit marked sensitivity to erastin, as evidenced by a change from the typical shape on stiff surfaces of spindle-like and spread to small and rounded, indicating cell death as observed under light microscopy (Figure 1B) (15). In contrast, cells seeded on soft hydrogels displayed no morphological changes, maintaining a small, rounded shape with and without erastin, as is typical for cells seeded on a soft surface (Figure 1B). To determine whether this effect was specific to erastin or a more general response to ferroptosis inducers, we treated all cell lines with two additional ferroptosis inducers of different mechanisms, FIN56 (class III) and RSL3 (class II), downstream effectors that inhibit the GPX4 enzyme (15, 31). We found a dose-dependent decrease in viability of MDA-MB-231 cells cultured on stiff hydrogels compared to those cultured on soft hydrogels, consistent across all three drugs (Figure 1, C-E). We further validated this finding across the three additional cell lines—CAOV3, RCC4, and HT1080—again demonstrating consistency across multiple cancer types (Supplemental Figure 1, A-I). Notably, we found that MDA-MB-231 and CAOV3 cell lines seeded on stiff hydrogels displayed higher cell viability after treatment with chemotherapy drugs carboplatin and taxol compared to those seeded on soft hydrogels, with the opposite trend observed for erastin. This supports the claim that stiff matrices selectively enhance susceptibility to ferroptosis induction while promoting relative resistance to conventional chemotherapeutic agents. (Supplemental Figure 3, A and B) (32).

To confirm that this difference in viability is due to ferroptosis rather than other types of cell death, we performed a rescue assay using ferrostatin-1, a ferroptosis inhibitor (33). Co-treatment with ferrostatin-1 and erastin fully rescued the viability of cancer cells on stiff hydrogels in all four cell lines, further supporting the role of ferroptosis (Figure 1F, Supplemental Figure 2, A-C). To further confirm the specificity of ferroptosis, we quantified lipid peroxidation, a key hallmark of ferroptotic cell death. Upon erastin treatment, cells cultured on stiff hydrogels showed markedly higher lipid peroxidation than those on soft substrates. This stiffness-dependent increase was observed in both MDA-MB-231 and HT1080 cells relative to their DMSO controls. (Figure 1, G and H). While cells displayed a degree of increased lipid peroxidation on soft matrices, the induction was dramatically enhanced for cells on stiff surfaces. Further, cells cultured on stiff hydrogels treated with erastin display increased expression of malondialdehyde, a lipid peroxidation byproduct, compared to cells cultured on soft hydrogels treated with erastin and DMSO controls (Figure 1I). Together, these results indicate that increased matrix stiffness enhances the sensitivity of cancer cells to ferroptosis-inducing agents, suggesting a mechanistic link between the tumor microenvironment and ferroptotic cell death.

### YAP regulates matrix-mediated ferroptosis

YAP is a transcriptional co-activator known to respond to matrix stiffness by translocating to the nucleus on stiff substrates (27). Additionally, these co-activators have been implicated in increasing sensitivity to ferroptosis (18, 26, 34). However, their role in regulating ferroptosis under matrix stiffness remains unclear. Consistent with previous literature, we found that all four cell lines, MDA-MB-231, HT1080, CAOV3, and RCC4 showed nuclear YAP localization when cultured on stiff hydrogels, whereas YAP remained cytoplasmic on soft hydrogels (Figure 2A, Supplemental Figure 4, A-C). This was further validated by increased mRNA expression of the YAP target gene *CTGF* (*CCN2)* in cells grown on stiff hydrogels compared to those grown on soft hydrogels (Figure 2B, Supplemental Figure 4D).

**Figure 2:**
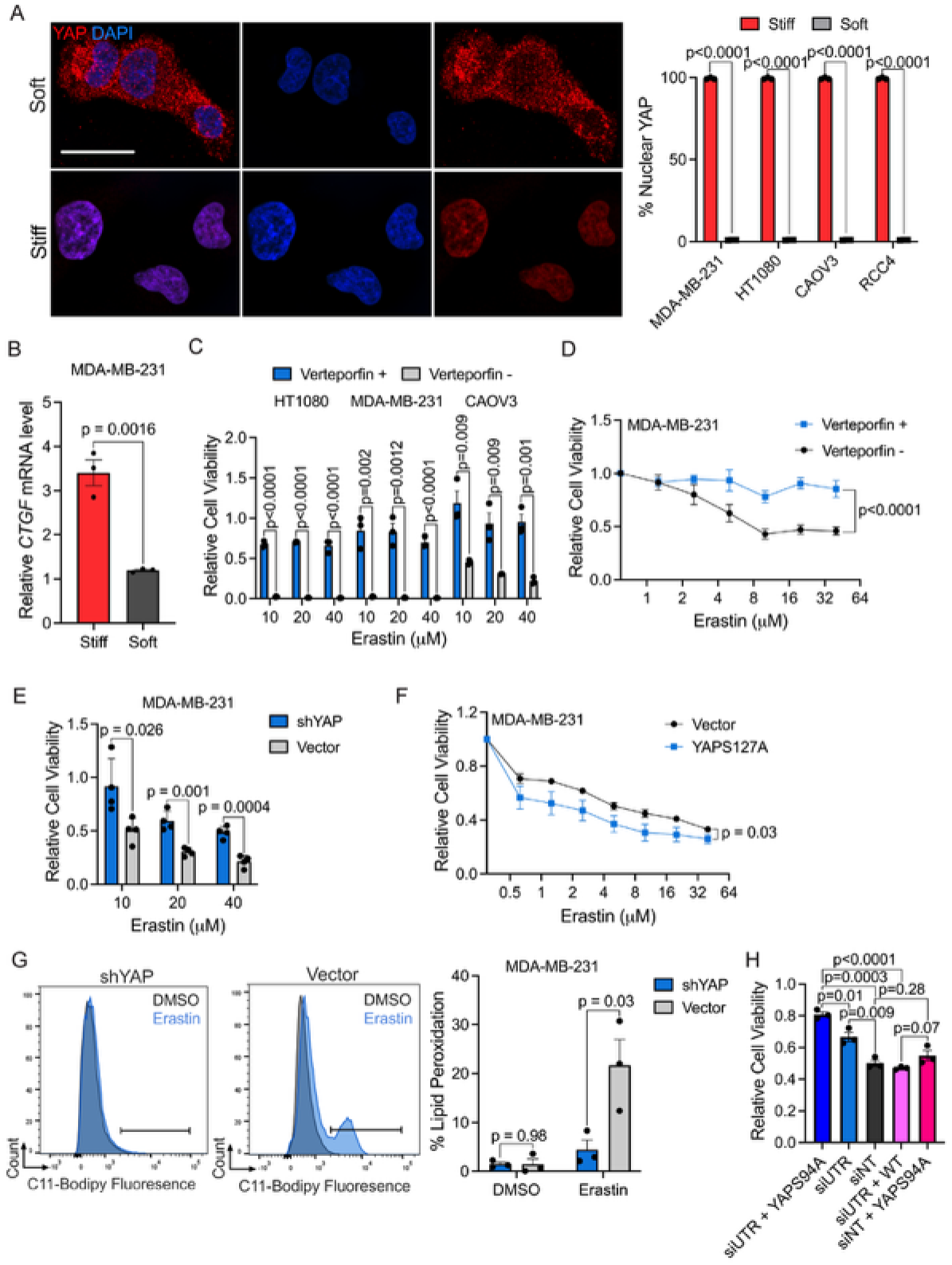
Matrix-mediated ferroptosis is regulated by YAP. (**A**) Representative immunofluorescence images (of N=3 independent experiments) MDA-MB-231 cells stained with DAPI (nuclei in blue) and YAP (in red) (left) and quantification for multiple cell lines (right; N=3) after 24hr in culture. Scale bars=25 μm (**B**) Relative mRNA expression of *CTGF* measured by RT-PCR of MDA-MB-231 cells seeded on soft or stiff hydrogels. (**C**) Relative cell viability of HT1080, MDA-MB-231 or CAOV3 cells on stiff hydrogels treated with erastin with or without 8 μM verteporfin (YAP inhibitor) for 16 hr. (N=3 for all cell lines) (**D**) Relative cell viability of MDA-MB-231 cells on soft hydrogels treated with erastin with or without 8 μM verteporfin (YAP inhibitor) for 16 hr. (**E**) Relative cell viability of shYAP MDA-MB-231 cells or vector treated with erastin for 16 hr. (N=3) (**F**) Relative cell viability of YAPS127A MDA-MB-231 cells or vector treated with erastin on soft hydrogels for 16 hr. (N=3) (**G**) Representative flow cytometry histograms of lipid peroxidation measured by C11-Bodipy fluorescence of either shYAP or vector MDA-MB-231 cell lines on stiff hydrogels treated with either 10 μM erastin or DMSO control for 6 hr (left) with quantification (right; N=3). (**H**) Relative cell viability of MDA-MB-231 cells treated with siNT or siUTR for *YAP1* rescued with YAPS94A mutant or WT YAP followed by 5 μM erastin treatment for 16 hr on stiff hydrogels. (N=3) All graphs represent means ± SEM. One-way ANOVA was used for A-C, E, G and H. Two-way repeated-measures ANOVA was used for D and F.

To assess the functional role of YAP in ferroptotic sensitivity, we cultured cells on stiff substrates and co-treated them with different doses of erastin and the YAP inhibitor, verteporfin (35). First, we confirmed that Verteporfin alone had no significant effects on cell viability (Supplemental Figure 4E) (35). We found that across three cell lines, verteporfin co-treatment led to a significant increase in cell viability compared to erastin alone (Figure 2C). This trend was also consistent for MDA-MB-231 cells cultured on soft substrates (Figure 2D). To further validate these findings, we used a previously established shYAP mutant MDA-MB-231 cell line and confirmed similar cell viabilities between shYAP and vector mutant cells (Supplemental Figure 4F) (34). Upon erastin treatment of cells cultured on stiff substrates, shYAP cells exhibited higher cell viability compared to the vector control, even at high erastin concentrations (Figure 2E, Supplemental Figure 4E). Compared to *YAP1* knockdown, siWWTR1 (TAZ), confirmed by western blot, did not alter the effects of erastin on cell viability (Supplemental Figure 5). Therefore, we concluded that YAP is the primary transcriptional co-activator mediating matrix-dependent ferroptosis in our system. Similar to MDA-MB-231, siRNA-mediated knockdown of *YAP1* in HT1080 cells led to significantly increased cell viability compared to the non-targeting control (Supplemental Figure 4, G and H). Reciprocally, YAPS127A mutant MDA-MB-231 cells (34) seeded on soft substrates showed decreased cell viability compared to vector control following erastin treatment (Figure 2F). To better understand the kinetics of *YAP1* knockdown in ferroptosis, we performed an EC50 time-course experiment. Cell viability began to decrease after 6 hours of erastin treatment for siNT-treated cells but remained high for siYAP-treated cells. Cell viability began to move closer between siYAP and siNT treated cells at 24 hours (Supplemental Figure 4I). Together, these results indicate that YAP activity enhances ferroptotic sensitivity and genetic or pharmacological inhibition of YAP confers resistance to erastin-induced cell death.

To confirm the specificity of this effect to ferroptosis, we measured lipid peroxidation levels of the shYAP mutant cell line. The shYAP MDA-MB-231 cell line displayed significantly reduced lipid peroxidation compared to the vector control (Figure 2G). Notably, lipid peroxidation levels in shYAP cells treated with erastin overlapped entirely with the DMSO control, whereas the vector control exhibited a distinct second population with high lipid peroxidation levels. (Figure 2G). Further, we used a YAPS94A mutant (mutation confirmed by Sanger sequencing) to determine whether our YAP-ferroptotic axis is TEAD dependent (Supplemental Figure 6A). To decrease WT YAP expression, we used a 3’UTR siRNA for *YAP1* or siNT control, then rescued with either the YAPS94A mutant, which cannot bind TEAD, or WT YAP. Decreased WT YAP activity, partial loss of function by YAPS94A, and the siUTR were confirmed by RT-PCR of *CTGF* (Supplemental Figure 6B). Further, restoration of YAP protein expression after addition of the WT YAP or YAPS94A were confirmed via western blot (Supplemental Figure 6C). Following erastin treatment, the siUTR and siUTR + YAPS94A mutants partially rescued cell viability, whereas siNT, siUTR + WT YAP, and siNT + YAPS94A mutant controls did not (Figure 2H). Therefore, matrix-dependent ferroptosis requires TEAD-dependent transcription. Together, these results suggest that YAP contributes to matrix-regulated ferroptosis sensitivity, where its inhibition confers resistance to ferroptosis.

### SKP2 mediates ferroptosis in cells cultured on stiff surfaces

Given the distinct second population observed in the lipid peroxidation data, we investigated differences in cell cycle regulation between stiffnesses following erastin treatment. Flow cytometry of cells stained with Propidium Iodide showed reduced accumulation of cells in the G2 phase when cultured on stiff surfaces compared to those cultured on soft surfaces (Figure 3A). This demonstrates that stiff surfaces accelerate cell cycle progression, resulting in less buildup in the G2 phase, consistent with previous literature linking stiffness to enhanced proliferation and cell cycle activity (1, 36, 37). Upon erastin treatment, cells cultured on soft surfaces displayed little to no change in G2 accumulation. Because of the slower cell cycle in soft surfaces, we suspect this dampens both proliferation and subsequent ferroptotic response. In contrast, cells cultured on stiff surfaces showed reduced G2 accumulation, highlighting a stiffness-specific interaction between cell cycle dynamics and ferroptosis (Figure 3A). To identify the molecular mediators linking these processes, we next examined YAP target genes involved in cell-cycle regulation. Among them, SKP2 —a key cell cycle driver and oncogene —emerged as a key candidate (38, 39). SKP2 is an F-box protein part of the SCF ubiquitin ligase complex, which targets proteins such as the tumor suppressor p27 for degradation to allow cells to pass through the G1-S checkpoint and progress in the cell cycle (38, 39). Overexpression of SKP2 is associated with various cancers and has been associated with resistance to chemotherapy in breast cancer cells (38, 40). Further, SKP2 has previously been shown to mediate ferroptosis, as *SKP2* knockdown decreased ferroptosis sensitivity (34).

**Figure 3:**
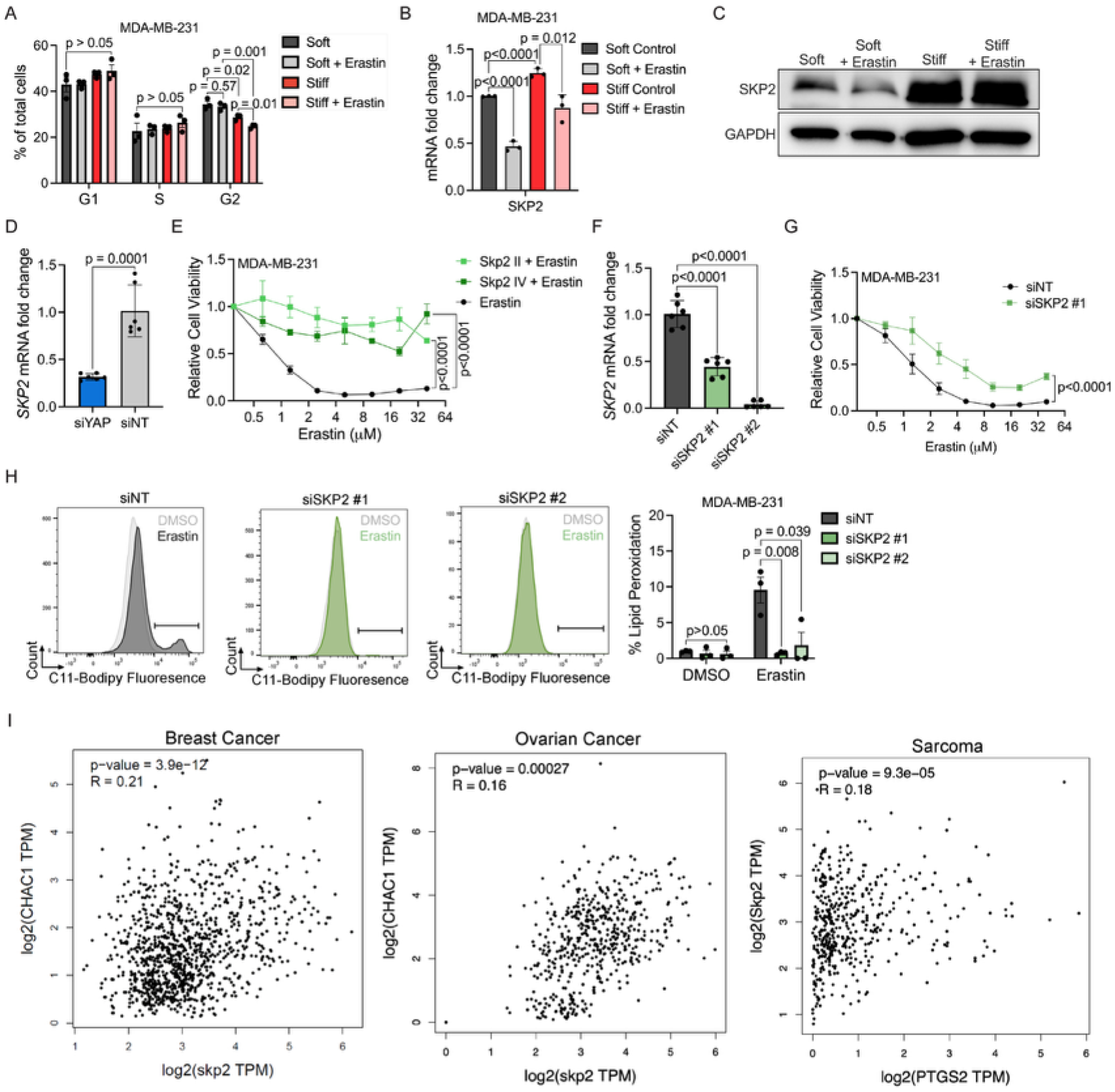
YAP target gene *SKP2* mediates ferroptosis. (**A**) PI flow cytometry of MDA-MB-231 cells treated with 10 μM erastin or DMSO on soft or stiff hydrogels for 6 hr. (**B**) mRNA fold change of *SKP2* by qRT-PCR of MDA-MB-231 cells treated with 10 μM erastin or DMSO for 8 hr on soft or stiff hydrogels. (**C**) Western blot of SKP2 and GAPDH. (**D**) mRNA fold change of *SKP2* by qRT-PCR of siYAP or siNT MDA-MB-231 cells. (n=6) (**E**) Relative viability of MDA-MB-231 cells on stiff hydrogels treated with erastin with or without 40 μM SKP2 inhibitor II or 10 μM SKP2 inhibitor IV for 16 hr. (**F**) mRNA fold change of *SKP2* by qRT-PCR of MDA-MB-231 cells transfected with siNT, siSKP2#1, or siSKP2#2 after 24 hrs. (n=6) (**G**) Relative viability of MDA-MB-231 cells on stiff hydrogels treated with erastin for 16 hr following siNT or siSKP2 #1 transfection. (**H**) *Left-*representative flow cytometry histograms of lipid peroxidation measured by C11-Bodipy of siNT, siSKP2 #1, or siSKP2 #2 transfected MDA-MB-231 cells on stiff hydrogels treated with 10 μM erastin or DMSO for 6 hr with *right-*quantification. (**I**) Human breast carcinoma, ovarian cancer, and sarcoma samples from the TCGA dataset plotted by GEPIA2 (43) for SKP2 correlation with *CHAC1* or *PTGS2*. (N=3 for all panels except I) All graphs represent means ± SEM. One-way ANOVA was used for A and B, D, F, and H. Two-way repeated-measures ANOVA was used for E and G.

In cells cultured on stiff surfaces, SKP2 was elevated on both the mRNA and protein levels compared to soft (Figure 3, B and C and Supplemental Figure 7, A and B). In stiff surfaces, erastin led to reduced *SKP2* mRNA levels, consistent with the previously observed decrease in G2 accumulation under ferroptotic stress (Figure 3B). Further, we confirmed that *YAP1* knockdown by siRNA reduced *SKP2* expression, as expected, validating its downstream regulation by YAP (Figure 3D). Further, verteporfin also led to decreased levels of *SKP2* in cells on stiff matrices but not in those on soft (Supplemental Figure 8). Next, we assessed whether SKP2 functionally mediates stiffness-dependent ferroptosis sensitivity. Co-treatment of erastin with either SKP2 inhibitor II (41) or IV (42) significantly increased cell viability compared to erastin alone in MDA-MB-231 breast cancer cells, HT1080 fibrosarcoma cells, and CAOV3 ovarian cancer cells (Figure 3E, Supplemental Figure 9, A and B). To further validate these findings, we performed *SKP2* knockdown using two independent siRNAs. RT-PCR confirmed reduced expression of *SKP2* by both siRNAs, achieving near-complete knockdown by siSKP2 #2 (Figure 3F). Following erastin exposure, *SKP2* knockdown improved cell viability on stiff surfaces relative to the siNT control (Figure 3G). To confirm the improved viability was due to reduced ferroptosis, we performed lipid peroxidation flow cytometry using both siSKP2 #1 and #2. Cells treated with siSKP2 displayed significantly lower lipid peroxidation levels, either entirely overlapping with the DMSO control or displaying a significantly diminished second population. In contrast, siNT cells showed a robust increase in lipid peroxidation, indicating greater ferroptotic stress (Figure 3H). Further, we compared erastin alone vs erastin combined with the SKP2 II inhibitor across a stiffness gradient from 0.1-100 kPa and tissue culture plastic (TCP). We found that SKP2 inhibition consistently rescued erastin-induced cell death across the stiffness spectrum, with the largest difference at stiffnesses above 8 kPa, supporting SKP2 as a downstream mediator of stiffness-dependent ferroptosis sensitivity (Supplemental Figure 10A). Given SKP2’s role in the cell cycle, we also assessed cell viability across multiple cell cycle stages. Following cell cycle synchronization in the MDA-MB-231 cells, we confirmed cell cycle stages using PI flow cytometry at 2 hr timepoints from 0-12hrs (Supplemental Figure 10, B and C). At each cell cycle stage we treated with either SKP2 II inhibitor, erastin, or combination and measured cell viability (Supplemental Figure 10D). The only significant difference in cell viability between erastin and the erastin combined with the SKP2 inhibitor was observed around the S phase cell cycle, when SKP2 activity is elevated (Supplemental Figure 10D). This provides additional evidence for the cell cycle-dependent role of SKP2 in matrix-dependent ferroptosis. Together, these findings demonstrate that stiffness-driven cell cycle dynamics contribute to ferroptosis sensitivity and identify SKP2 as a critical mediator whose inhibition drastically dampens ferroptosis sensitivity on stiff surfaces.

Analysis of human patient data from the TCGA dataset (43) revealed that *SKP2* expression correlates with distinct ferroptosis markers across tumor types—*CHAC1* in breast and ovarian cancers, and *PTGS2* in sarcomas. Although correlations between *SKP2* and individual ferroptosis-associated genes were modest, several were statistically significant, suggesting that *SKP2* expression is associated with ferroptosis-related transcriptional features in patient tumors (Figure 3I). Moreover, survival analysis of human patients from either Kaplan-Meier Plotter (44) or Xena databases (45), shows that high *SKP2* expression in tumors correlates with poorer survival outcomes in human renal carcinoma, sarcoma, breast cancer, and ovarian cancer (Supplemental Figure 11, A-D). Next, we compared survival curves for breast cancer in low vs high *SKP2* and low vs high *CHAC1* cohorts (44). Stratified analyses showed that high *SKP2* remained associated with poor recurrence-free survival in both *CHAC1*-high and *CHAC1*-low tumors. Likewise, high *CHAC1* was associated with poorer survival in both *SKP2*-high and *SKP2*-low cohorts. Consistent with our proposed model, comparison of the survival estimates found that the *SKP2*-high/*CHAC1*-high subgroup exhibited the poorest outcome, whereas the *SKP2*-low/*CHAC1*-low subgroup exhibited the most favorable outcome (Supplemental Figure 11, E and F). Together, these analyses support the potential clinical relevance of the YAP-SKP2 axis across the tumor types examined (Figure 4).

**Figure 4:**
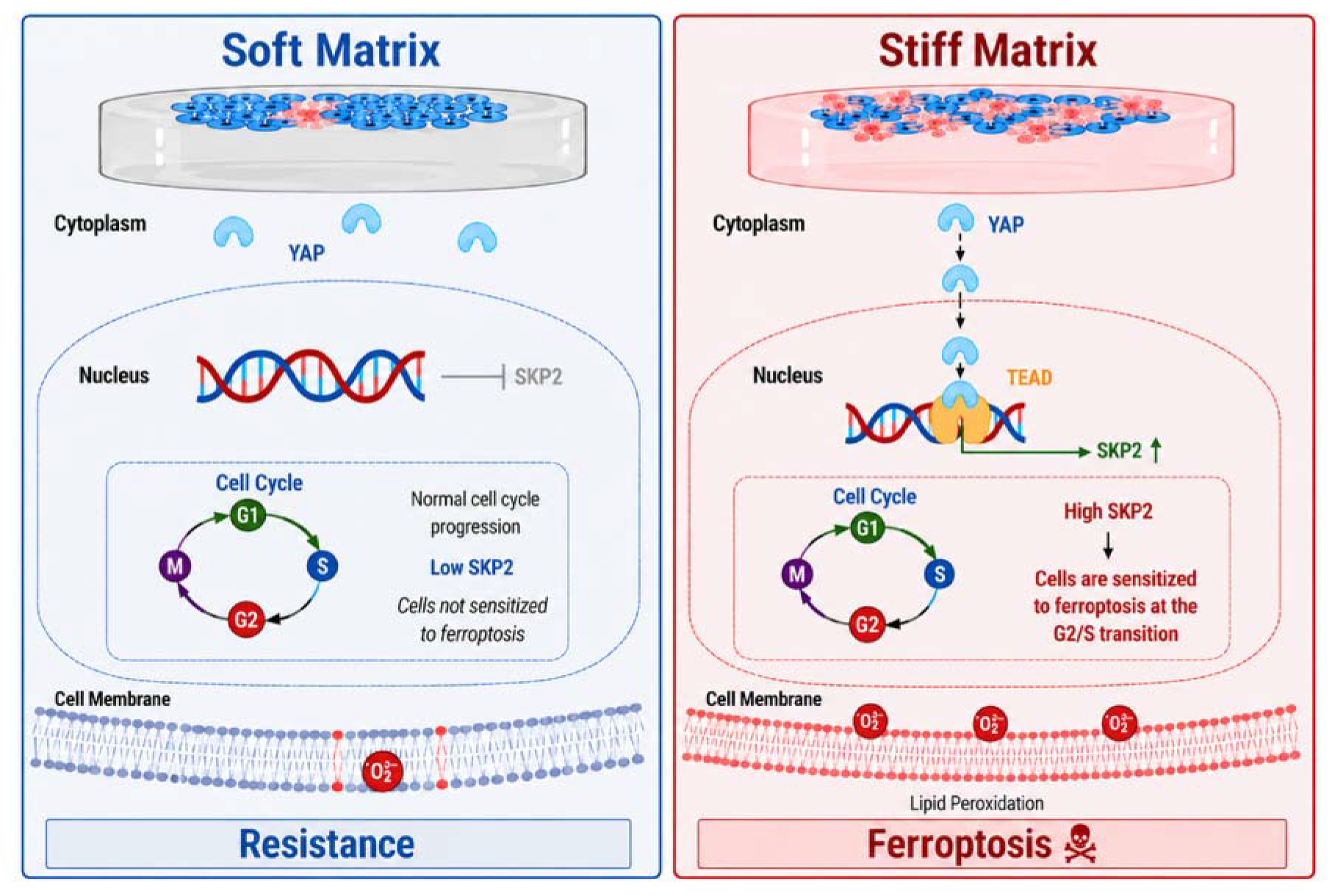
Matrix-mediated ferroptosis is regulated by a YAP-SKP2 signaling axis.

### Ferroptosis can target stiff chemoresistant tumors

After establishing that matrix stiffness regulates ferroptosis in vitro, we sought to examine whether this pathway could be leveraged to treat stiff, chemoresistant tumors in vivo. To generate stiffened tumors, we encapsulated MDA-MB-231 breast cancer cells in a semi-synthetic gelatin and hyaluronan-based injectable hydrogel (46, 47). Under anesthesia, the hydrogel-cell mixture was subcutaneously injected into the right rear flank of immunodeficient NOD/SCID female mice. Tumors developed for seven days before initiating treatment to reach a size of 50-150 mm^3^ (Figure 5A).

**Figure 5:**
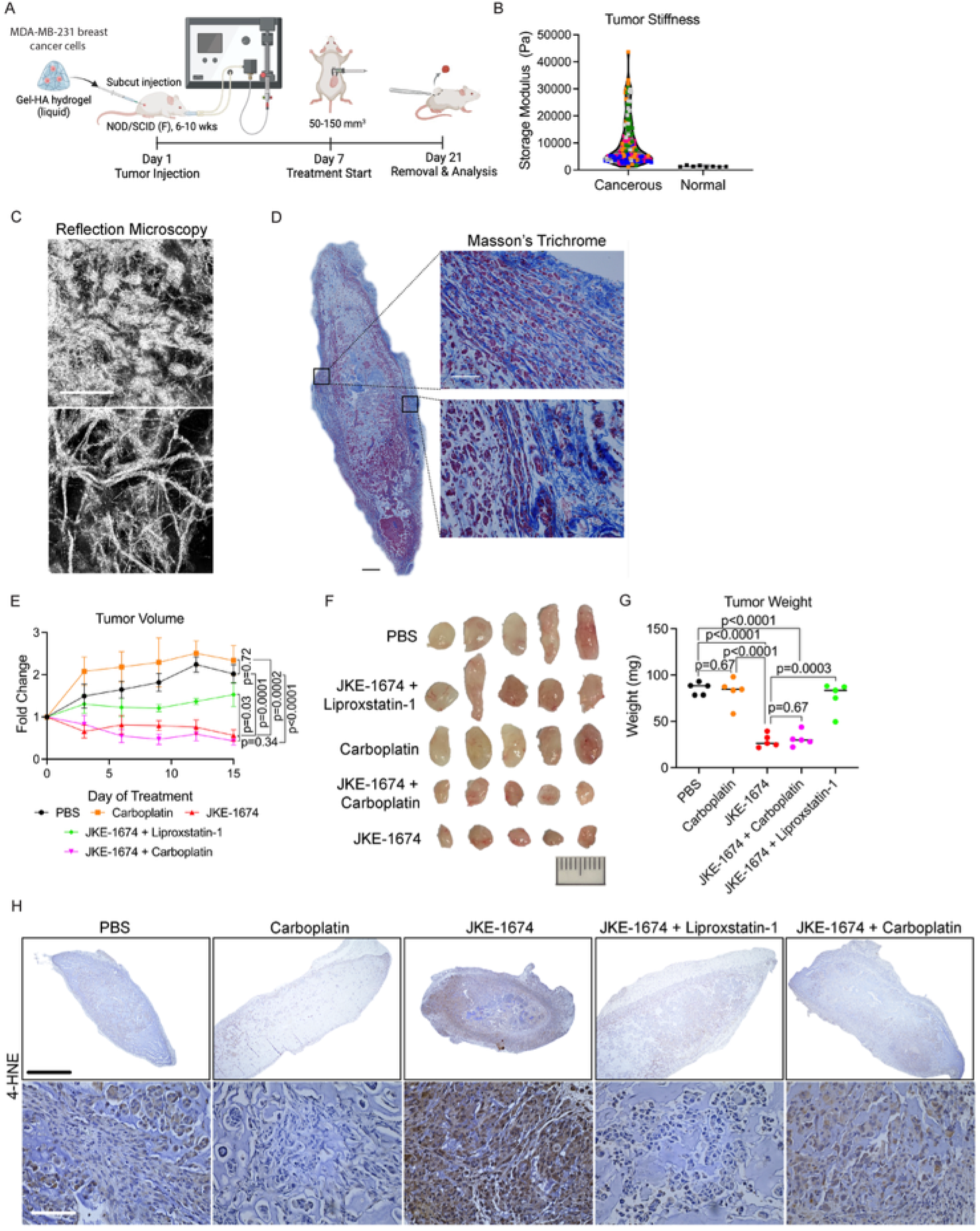
Ferroptosis is a potential therapeutic option for stiffened chemo-resistant breast tumors. (A) Schematic of experimental timeline for establishing stiffened tumors in vivo. (B) AFM measurements of tumor sections at 10 Hz for 5 individually isolated tumors and normal human breast tissue from (29). (C) Collagen reflection microscopy of isolated tumor sections treated with PBS. Scale bars=100 μm (D) Masson’s trichrome staining of isolated tumor sections treated with PBS. Scale bars=50 μm (E) Fold change in tumor volume, measured by calipers, relative to the initial tumor volume on day 7 after treatment initiation. (F) Images of isolated tumors for each condition. Scale bar = 10 mm (G) Measured weight of whole isolated tumors for each condition. (H) 4-HNE histological staining of each condition. Top scale bar=1000 μm. Bottom scale bar=100 μM (N=5 for all conditions and measurements). Representative images are from tumors collected from N = 5 mice per group. All graphs represent means ± SEM except for B and G. Lines in graph G represent mean. One-way ANOVA was used for G. Planned pairwise comparisons of longitudinal tumor-growth trajectories were analyzed using two-way repeated-measures ANOVA; P values for the indicated treatment comparisons are shown in the figure.

To directly assess tumor stiffness, we performed atomic force microscopy (AFM) on fresh tumor sections, which revealed a high average storage modulus and heterogeneous mechanical properties ranging from 1–43 kPa, values and ranges higher than normal physiological breast tissue (0.5–1 kPa) (Figure 5B, Supplemental Figure 12) (6, 29). Accordingly, we interpret these tumors as relatively stiffened rather than as uniformly reaching the 25 kPa stiffness used in our in vitro studies. The observed spatial heterogeneity in solid tumors is expected due to regional differences in cellularity, matrix deposition, vascularization, edema, and necrosis, with the same tumors containing localized regions of markedly elevated stiffness (6, 30). We also assessed ECM composition and structure using Masson’s trichrome staining and reflection microscopy. We found fibrillar collagen deposition and a heterogeneous ECM structure, consistent with the well-documented properties of solid tumors (Figure 5, C and D).

Mice were then randomly assigned to one of five treatment groups: 1) PBS control, 2) carboplatin (chemotherapy), 3) JKE-1674 (ferroptosis inducer), 4) JKE-1674 + liproxstatin-1 (ferroptosis inhibitor, to distinguish ferroptosis from apoptosis), and 5) JKE-1674 + carboplatin (combination therapy) (48, 49). Carboplatin was selected for its clinical relevance in treating triple-negative breast cancer and for alignment with our chosen cell line. (50) JKE-1674 was selected because it is a GPX4-targeting ferroptosis inducer that has been used to trigger ferroptosis in vivo (51). Tumor volume was measured every two days and normalized to the initial measurement on day 7 post-injection (the first day of treatment). Consistent with the expected chemoresistance of stiffened tumors (9, 32), no significant differences in tumor growth were observed between carboplatin-treated and PBS-treated mice (Figure 5E). In contrast, JKE-1674 treatment resulted in a marked reduction in tumor growth compared to both PBS and carboplatin groups (Figure 5E). Importantly, co-treatment with JKE-1674 and liproxstatin-1 rescued tumor growth, confirming that the reduction in tumor volume in the JKE-1674 treated group is due to ferroptosis rather than apoptosis or other forms of cell death (Figure 5E). The combination of JKE-1674 and carboplatin showed an insignificant reduction in tumor growth relative to JKE-1674 alone (Figure 5E). Representative images of excised tumors further supported these findings where PBS-, carboplatin-, and JKE-1674 + liproxstatin-1-treated mice showed similar tumor size and JKE-1674- and JKE-1674 + carboplatin–treated mice had significantly smaller tumors (Figure 5F). These trends were also consistent for final tumor weight (Figure 5G). Immuno-staining shows an elevated degree of 4-HNE, a lipid peroxidation byproduct, in JKE-1674-treated tumors compared to all other treatment groups (Figure 5H).

To further validate this result, a counterpart of soft tumors was generated with the same experimental setup but with a hydrogel composition tuned to form a softer matrix. Tumor volume curves showed no significant differences between soft tumors treated with PBS and those treated with JKE-1674 (Supplemental Figure 13B). Further, images of the tumor reveal tumors of similar size (Supplemental Figure 13C). Lastly, the trend stays consistent for tumor weight as well, with no significant differences (Supplemental Figure 13D). YAP stains and quantification for percent of nuclear localization showed significantly higher levels of nuclear YAP in stiffened tumors compared to soft tumors (Supplemental Figure 14A). Therefore, nuclear YAP levels correlated strongly with the response to ferroptosis induction in vivo. Moreover, SKP2 staining showed overall higher levels and greater nuclear overlap in stiffened tumors compared to soft tumors, validating our in vitro findings in our tumor model (Supplemental Figure 14B).

To assess whether this effect was consistent for other tumor types, we next examined CAOV3 ovarian cancer cells under the same experimental conditions, comparing PBS and JKE-1674 (Supplemental Figure 15A). Here too, CAOV3 tumors exhibited increased stiffness as measured by AFM compared to normal ovarian tissue (52) (Supplemental Figure 15B) and high, heterogeneous ECM deposition by Masson’s trichrome staining (Supplemental Figure 15C). Consistent with the trend in MDA-MB-231 tumors, JKE-1674 treatment resulted in little to no tumor growth over time, whereas PBS-treated tumors displayed a steady or exponential increase in volume (Supplemental Figure 15D). Together, these data demonstrate that ferroptosis induction is an effective treatment for targeting stiff, chemoresistant tumors.

## Discussion

Here, we identify matrix stiffness as a key regulator of ferroptosis through the YAP–SKP2 axis. This relationship was consistently reproduced across four distinct cancer cell types—breast, ovarian, renal, and fibrosarcoma—representing both epithelial and mesenchymal lineages. The reproducibility across four cancer cell types suggests that stiffness-regulated ferroptosis is not restricted to a single tumor lineage and may have broader relevance across cancers. Using soft and stiff PA hydrogels, we showed across four tumor types that cells cultured on stiff substrates exhibit higher sensitivity to ferroptosis through lower cell viability and higher lipid peroxidation compared to those on soft. Confirming this trend across four tumor types suggests that matrix-regulated ferroptosis is relevant across multiple tumor types and may be generally important for other organ systems not explored here. YAP has been shown to shuttle to the nucleus and influence gene expression on stiff substrates (27). Nuclear YAP under high cell density was reported to increase sensitivity to ferroptosis (26). Consistent with these findings, we showed that this matrix-dependent ferroptosis sensitivity was regulated by nuclear YAP, as demonstrated by inhibition and activation studies. Knockdown of YAP via drug and genetic inhibition rescued ferroptosis shown through higher cell viability and decreased lipid peroxidation. Overactivation through the YAPS127A mutant protein led to lower cell viability. This provides additional context for the regulation of ferroptosis by matrix stiffness via YAP.

After investigating differences in cell cycle dynamics, we identified SKP2 as a central regulator of matrix-dependent ferroptosis. Prior studies have established that YAP/TAZ activity regulates ferroptosis susceptibility, including through SKP2, with cell density modulating Hippo signaling (26, 34). Building on this framework, our work identifies extracellular matrix stiffness—the mechanical cue most characteristic of solid tumors—as an upstream regulator of ferroptosis susceptibility. Notably, although matrix stiffness confers resistance to conventional chemotherapy, it concurrently sensitizes tumor cells to ferroptosis, revealing a mechanically defined therapeutic vulnerability that can be exploited to regress stiff, treatment-refractory tumors in vivo. Therefore, building on previous results of SKP2 regulation in cell-density dependent ferroptosis, drug and genetic inhibition of SKP2 displayed increased cell viability and decreased lipid peroxidation in matrix-mediated ferroptosis (4, 5, 34). Further, *SKP2* correlated with expression of ferroptosis markers and lower overall survival in human patient data. Since SKP2 expression has been proposed as a tumor biomarker, SKP2 may serve as a mechanically informed biomarker for identifying stiff, YAP-active tumors likely to respond to ferroptosis-based therapies. However, because SKP2 inhibition suppresses ferroptotic sensitivity, SKP2-targeting agents should not be combined concurrently with ferroptosis inducers, as such combinations may diminish therapeutic efficacy.

It has been shown extensively that matrix stiffness leads to therapeutic resistance (3–5). Previously, semi-synthetic gelatin-hyaluronan hydrogels have been used to model stiff subcutaneous tumors (46, 47). Building on this system, we established a stiff tumor xenograft model and found that these stiffened tumors were vulnerable to the ferroptosis-inducer JKE-1674 but resistant to chemotherapy, as demonstrated by decreased tumor volume and weight. Therefore, this provides evidence for ferroptosis as a therapeutic approach for stiff, treatment-refractory tumors. Further, these findings demonstrate the potential translatability of exploiting mechanotransduction pathways to overcome stiffness-associated drug resistance through a metabolic predisposition to ferroptosis. Importantly, our engineered tumors aim to replicate a stiff and mechanically heterogeneous tumor ECM, rather than a tissue with uniformly high modulus, since achieving a precisely controlled and spatially uniform tissue stiffness in vivo continues to be a challenge. Importantly, because the soft and stiffened tumor models differ in hydrogel composition, and direct mechanical characterization of the soft tumors was not performed, we cannot attribute the observed in vivo differences exclusively to matrix stiffness, and contributions from compositional differences between the hydrogel formulations cannot be excluded. However, matrix stiffness has also been suggested to protect cells from ferroptosis in other studies. In hepatocellular carcinoma (HCC), matrix stiffness induced PD-L2, which functioned as a suppressor of xCT-mediated ferroptosis and led to sorafenib resistance and increased cancer growth (53). This highlights the importance of context in matrix-dependent ferroptosis sensitivity. Future studies should screen across varying stiffnesses, tumor types, and mechanisms of ferroptosis induction, which may influence sensitivity or resistance to ferroptosis. These findings will be essential to inform the potential use of ferroptosis as part of future clinical regimens for tumors. Looking at our chosen in vitro model, PA gels are the most established platform to study ECM stiffness in 2D. While reductionist, PA gels offer a powerful model to reveal fundamental links. Although we validated our results with a 3D in vivo hydrogel, future research should utilize 3D in vitro hydrogel models to explore how additional mechanical properties like viscoelasticity and plasticity influence ferroptosis sensitivity (54, 55).

In our xenograft model, a hydrogel system was employed to ensure the presence of a defined stiff matrix within the tumor microenvironment. This approach necessitated implantation of isolated tumor cells, which in turn required the use of immunodeficient mice. While this model provides important insight, future studies should incorporate stiffened tumor models in immunocompetent mice to better capture the complexity and clinical relevance of tumor–immune interactions. Notably, the stiffness-dependent enhancement of ferroptosis was consistent across collagen- and gelatin and hyaluronic acid–based hydrogels, indicating the effect is driven by mechanics rather than a specific matrix composition. However, the tumor microenvironment is considerably more complex, and future studies should investigate how additional ECM components, such as fibronectin and laminin, influence ferroptosis sensitivity and the YAP–SKP2 signaling axis. Likewise, stromal and immune cells play important roles in ECM remodeling and tumor progression, and their interactions with tumor cells may further modulate stiffness-dependent ferroptosis (56, 57). Incorporating these components into future models will be important for understanding how the broader tumor microenvironment influences ferroptosis and its therapeutic potential. Although the mechanistic studies presented here were performed primarily in established cancer cell lines, analyses of TCGA and KM Plotter datasets further support the clinical relevance of SKP2 in human tumors. Nevertheless, established cell lines cannot fully recapitulate the cellular heterogeneity, ECM architecture, and biomechanical complexity of patient tumors. Future studies using primary human specimens, patient-derived organoids, and patient-derived xenograft models will be important to determine whether the stiffness-dependent YAP–SKP2–ferroptosis axis identified here is similarly conserved in human disease.

The use of biomaterials and complex microphysiological systems is currently limited in the context of ferroptosis. Our results showing matrix-dependent ferroptosis sensitivity indicate a strong need to implement these systems and other mechanical forces in the future. These results can set the foundation for the use of engineering principles in studying ferroptosis and the potential for ferroptosis-inducers as a treatment route for stiff tumors. Importantly, our findings do not imply that tumor stiffness must be mechanically reversed to achieve therapeutic benefit. Rather, stiffness establishes a YAP–SKP2 transcriptional state that predisposes tumor cells to ferroptosis. Therapeutic induction of ferroptosis exploits this stiffness-imposed signaling landscape, allowing selective targeting of stiffened tumors without altering the mechanical microenvironment itself. Collectively, our work establishes stiffness-driven ferroptosis as a therapeutically exploitable dimension of mechanotransduction and positions the YAP-SKP2 axis as a tractable vulnerability in stiff, treatment-refractory cancers.

## Materials & Methods

### Sex as a biological variable

Our study exclusively examined female mice because the disease modeled is only relevant in females.

### Study Design

This study was designed to test the pre-specified hypothesis that matrix stiffness is an exploitable vulnerability to ferroptosis in stiff treatment-refractory tumors. Multiple human cancer cell lines were cultured on 2D polyacrylamide soft and stiff hydrogels or embedded in an in vivo injectable hydrogel in a murine xenograft model. Three to four biological replicates, independent cultures established on different days, were chosen as the sample size for in vitro studies and three to five were chosen for in vivo studies. Data collection endpoints were prospectively defined at 16 hours for cell viability and 6 hours for lipid peroxidation studies. Experiments were excluded only if hydrogel functionalization with collagen protein failed defined as poor adhesion following cell seeding or cultures exhibited contamination. Studies were performed as a controlled laboratory experiment and with measurement technique details outlined below. Hydrogel stiffness (soft vs. stiff) or pharmacologic treatments (i.e. ferroptosis induction, inhibitors, or vehicle, etc.) were applied in a fully crossed design. For mouse experiments, following tumor implantation, mice were randomized into treatment conditions to prevent bias but treatments were not performed blindly. Further only female mice were used given the types of cancer were female breast cancer and ovarian cancer, only applicable to female populations.

### Polyacrylamide hydrogels

Soft and stiff polyacrylamide wellplates were fabricated as described previously (58) or acquired from Matrigen (SW6-EC-0.5-EA, SW6-EC-25-EA, SW96-EC-0.5-EA, or SW96-EC-25-EA), coated with rat-tail collagen I for 1 hr, and washed twice with ice-cold PBS prior to cell culture.

### Cells

HT1080 (ATCC), MDA-MB-231 (ATCC), RCC4 (originally gifted by Dr. Denise Chan (Department of Radiation Oncology, University of California, San Francisco CA), and CAOV3 (originally gifted by Dr. Zhiqing Huang and Dr. Susan Murphy, Duke University, Durham NC) cells were cultured in DMEM (GIBCO) supplemented with 10% heat-inactivated fetal bovine serum (Cytiva) and 1% penicillin-streptomycin (Thermo Fisher Scientific) in a humidified incubator at 37°C and 5% CO2. shYAP expressing, YAPS127A, and vector MDA-MB-231 cells were generated as previously described (34).

### Cell Viability

Cell viability was assessed using the CellTiterGlo luminescent cell viability assay kit (Promega) following the manufacturer’s protocol. Cell Titer is added to the media, shaken for 2 min, and incubated for 10 min before reading luminescence on the plate reader.

### C11 Bodipy Lipid peroxidation flow cytometry

Lipid ROS levels were measured by 1.5 μM of C11-BODIPY dye (D3861, ThermoFisher Scientific) after treatment with 6 hr erastin or DMSO control. The culture media was replaced with 1.5 μM of C11-BODIPY dye for 30 min. After, the cells were harvested by TrypLE and washed three times with ice-cold PBS, followed by resuspending them in PBS plus 1% BSA. The amount of ROS within cells was examined by flow cytometry analysis (FACSCantoTM II, BD Biosciences).

### Western blot analysis and quantification

For immunoblotting, cells were washed with ice-cold phosphate-buffered saline (PBS), lysed in RIPA buffer (Sigma), supplemented with Halt™ Protease and Phosphatase Inhibitor Cocktail (ThermoFisher Scientific). Proteins were quantified by BCA protein assay (ThermoFisher Scientific). Equal amounts of proteins were resolved by sodium dodecyl sulfate–polyacrylamide gel electrophoresis (SDS-PAGE) and transferred to the polyvinylidene difluoride (PVDF) membrane (Millipore). The membranes were blocked with 5% non-fat milk and then probed with YAP (Santa Cruz, #sc-271134) TAZ (Cell signaling, #70148), β-actin (CST, #3700S), SKP2 (Thermo Fisher Scientific, #32-3300), GAPDH (CST, #5174T), or β-Tubulin (CST, #2141) followed by HRP-conjugated secondary antibodies (Rabbit: CST #7074S, Mouse: CST #7076S). The immunosignals were achieved by the Pierce™ SuperSignal™ West Pico PLUS Chemiluminescent Substrate western blotting detection agent (ThermoFisher Scientific) and detected by a Bio-Rad ChemiDocTM Imaging System. Blots were analyzed using ImageJ, and bands were normalized to β-actin or GAPDH expression.

### Immunofluorescence staining

Cells were cultured on polyacrylamide hydrogels to an appropriate density. Cells were fixed with 4% formaldehyde for 10 min and then washed three times with PBS. After blocking in 1% BSA and permeabilized with 0.1% Triton-X (Sigma-Aldrich) for 20 min, gels were incubated with the YAP antibody (1:200; SantaCruz, #sc-15407) diluted in appropriate antibody diluent (ThermoFisher Scientific) overnight. After washing with 0.05% Tween-20 in PBS, gels were incubated with Alexa Fluor 546-conjugated secondary antibodies (1:400; Invitrogen, #A11035) for 1 hr, and DAPI (Invitrogen, #D1306) for 10 min. For malondialdehyde, cells were incubated with the antibody stain (1:200, Invitrogen, #MA5-45805) for 2 hours. Images were acquired using a Nikon confocal microscope equipped with a 60X objective.

### RNA isolation and quantitative real-time PCR

RNA was extracted using TRIzol reagent (Invitrogen) and purified by RNeasy mini kit (Qiagen). Complementary DNA was synthesized using GoScript reverse transcription mix (Promega). Gene expression was performed by real-time RT-PCR, using SybrGreen primers and mastermix (Thermo Fisher Scientific). *TBP* served as the housekeeping gene. The ΔΔCt method was used for data analysis. Primer sequences were as followed: *TBP*, Fw: GCC AAG AGT GAA GAA CAG and Rev: GAA GTC CAA GAA CTT AGC TG; *SKP2*, Fw: ATGCCCCAATCTTGTCCATCT and Rev: CACCGACTGAGTGATAGGTGT; *CTGF*, Fw: GCCAATGAGCTGAATGGAGT and Rev: CAATCCGGTGTGAGTTGATG.

### siRNA transfection

Cells were transfected with SMARTpool:siGENOME and Non-Targeting Control siRNA #1 according to the manufacturer’s protocol. Briefly, cells were seeded on a six-well plate overnight in antibiotic-free media and treated with 50 nM siRNA for 48 h. The knockdown was confirmed via quantitative real-time fluorescence polymerase chain reaction (qRT-PCR) or western blot after 24 h.

### Cell Cycle Propidium Iodide flow cytometry

Cells were isolated using TrypLE, spun down at 400g for 5 min, and fixed in 70% ice-cold sterile ethanol and left at 4 °C overnight. The following day, the fixed cell solution was spun down at 10,000g for 5 mins and the supernatant ethanol was removed by aspirating. The cell pellet for each condition was resuspended in 50 µL RNAse (10 µg µL^−1^) to reduce background stain. Following this, 425 µL of cell staining buffer (Biolegend) and 25 µL of PI staining reagent (Biolegend, #421301) was added to the cell pellet and incubated for 45 min at RT in the dark. The stained cells were analyzed on a BD FACSCanto Flow cytometer for cell cycle at the Duke Cancer Institute Flow Cytometry core facility. The data were analyzed using the FlowJo software (Ashland, OR) and peaks were identified as G1 (2N), S, or G2 (4N) phase.

### Cell Synchronization

MDA-MB-231 cells were synchronized using a double-thymidine block. Cells were seeded in 6-well plates and grown to approximately 40% confluence before treatment with 2 mM thymidine for 14 h. Cells were then washed twice with PBS and released into complete DMEM supplemented with 24 μM deoxycytidine for 9 h, followed by a second 14-h treatment with 2 mM thymidine. After the second block, cells were washed twice with PBS and released into complete DMEM containing 24 μM deoxycytidine. Cells were collected every 2 h from 0 to 12 h after release, and cell-cycle distribution was assessed by propidium iodide flow cytometry. At each time point, parallel cultures were treated with 10 μM erastin for 6 h, after which cell viability was measured using the CellTiter-Glo assay.

### EC50 time course experiment

MDA-MB-231 cells were transfected with siYAP or siNT then treated with 10 μM erastin and measured for cell viability by CellTiter-Glo at hours 0, 2, 4, 6, 8, 12, 16 and 24. Cell viability for each were normalized to DMSO controls for siYAP and siNT respectively at each time point.

### YAPS94A mutant generation

The YAP S94A expression plasmid was obtained from Addgene (plasmid #59145), propagated in bacterial cells under appropriate antibiotic selection, and purified using standard plasmid isolation procedures (59). The plasmid sequence was verified by Eton Bioscience, and no unintended mutations were detected. MDA-MB-231 cells were seeded in 6-well plates at 1.65 × 10^5 cells per well and transfected with *YAP1* 3′UTR-targeting siRNA (si-3′UTR) or non-targeting control siRNA (siNT) (Horizon Discovery) for 24 h, as described above. The medium was then replaced, and the cells were allowed to recover for 6 h before plasmid transfection. For each well, 7 µL Lipofectamine 3000 (Thermo Scientific) was diluted in 125 µL Opti-MEM, while 1 µg of either YAP S94A or wild-type YAP plasmid DNA was combined with 3 µL P3000 reagent (Thermo Scientific) in a separate 125 µL volume of Opti-MEM. The two solutions were combined, incubated for 10 min at room temperature, and added to cells containing 2 mL culture medium. After 10 h, the transfection medium was replaced with complete DMEM, and cells were incubated overnight. Cells were subsequently reseeded, treated with erastin for 16 h, and cell viability was measured using the CellTiter-Glo assay.

### Data mining

Human breast carcinoma, ovarian cancer, and sarcoma samples from the TCGA dataset were plotted by GEPIA2 (43) for pair-wise gene expression correlation analysis with Pearson correlation coefficient in Figure 3. Human breast and ovarian cancer samples were plotted for Kaplan-Meier survival plots by Kaplan-Meier Plotter (44). Human renal carcinoma and sarcoma samples from the TCGA dataset were plotted for Kaplan-Meier survival plots by UCSC Xena (45). For all Kaplan-Meier analyses, patients were dichotomized into high- and low-expression cohorts at the median expression value of the gene of interest; the auto-select-best-cutoff option was not used. This yielded cohorts of essentially equal size (e.g., n = 1,239 and n = 1,221 for the breast cancer analysis), with minor departures from an exact split reflecting patients sharing the identical expression value at the median. Hazard ratios with 95% confidence intervals and log-rank *P* values were computed by the tool. For the combined analyses in Supplemental Figure 11 E and F, the cohort was first restricted by [*SKP2* or *CHAC1*] expression relative to the median and then stratified at the median expression of the second gene.

### YAP and SKP2 image analysis

Nuclear fluorescence enrichment was quantified per z-plane on the raw confocal stacks. Nuclei were segmented using the DAPI channel, and the mean YAP (red channel) or SKP2 (green channel) fluorescence intensity within the nuclear region was divided by the mean fluorescence intensity in a thin perinuclear region, generating a nuclear enrichment index. Therefore, giving a ratio where >1 indicates nuclear enrichment and <1 nuclear exclusion. Values were aggregated to one per animal (N = 5 per group).

### Animal Study

#### Study Approval

All animal care and experiment procedures were conducted according to the National Institutes of Health guidelines. The animal protocols were approved by the Division of Laboratory Animal Resources of Duke University (Protocol #A009-24-01).

#### Tumor Study

NOD/SCID female mice of age 6-10 weeks (Jackson Laboratory) were anesthetized with isoflurane and injected subcutaneously with 100 uL of semi-synthetic hyaluronan-derived gel precursor solution and 2 million MDA-MB-231 cells or CAOV3 cells. Mice remained anesthetized for 20 min after injection to undergo gelation. Tumors were allowed to form over 7 days for MDA-MB-231 cells and 5 days for CAOV3 cells before the start of treatment. Mice were randomized and labeled by ear punch into 5 groups: PBS, 80 mg/kg weekly carboplatin (MedChemExpress), 25 mg/kg JKE-1674 daily (Selleckchem), 25 mg/kg JKE-1674 + 10 mg/kg Liproxstatin-1 daily (MedChemExpress), 25 mg/kg JKE-1674 daily + 80 mg/kg weekly carboplatin. To monitor tumor growth, tumors were measured by caliper every 2 days. Mice were euthanized at the humane or experimental endpoint (21 days after tumor injection). Tumors were resected, imaged, measured, and weighed.

#### Semi-synthetic hyaluronan-derived gels

For stiffened tumors, hydrogels were fabricated as previously described (46, 47). In brief, the hystem-C kit (Advanced Biomatrix), 5 mg/mL Glycosil (thiol-modified hyaluronan), 10 mg/mL Gelin-S (thiol-modified gelatin), and 1.5 mg/mL Extralink (PEGDA) were combined at ratios of 2:2:1 respectively. For soft tumors, the same kit was used but the formulation was adjusted to establish a soft matrix. 2.5 mg/mL Glycosil, 10 mg/mL Gelin-S, and 0.3 mg/mL Extralink were combined at ratios of 2:2:1, respectively. The gel precursor mixture is combined with tumor cells and injected into NOD/SCID mice.

#### Histology

Tumors were submerged in 4% formaldehyde for 24 hrs at 4°C then replaced with 70% ethanol. Paraffin embedding and sectioning were performed by the Duke Pathology and Histology core. Masson’s trichrome were performed by the Duke Pathology and Histology core. For 4HNE-staining, paraffin-embedded tissue sections were heated at 65°C for 30 min, deparaffinized in xylene, and rehydrated through a graded ethanol series to distilled water and PBS. Antigen retrieval was performed in citrate buffer (10 mM sodium citrate, 0.05% Tween-20, pH 6.0) at 100°C for 5 min. Sections were permeabilized with 0.5% BSA and 0.4% Triton X-100 in PBS, and endogenous peroxidase activity was blocked using 1% hydrogen peroxide. Following blocking in normal serum diluted in 1% BSA and 0.4% Triton X-100, sections were incubated overnight at 4°C with an anti-4-hydroxynonenal (4-HNE) primary antibody (Bioss Antibodies, #BS-6313R; 1:500). Sections were subsequently incubated with a biotinylated secondary antibody (Vector Laboratories, #BA-1000-1.5) for 2 h at room temperature, followed by ABC reagent (Vector Laboratories) for 1–1.5 h. Immunoreactivity was visualized using ImmPACT DAB substrate (Vector Laboratories), with development monitored microscopically and terminated by immersion in distilled water. Sections were counterstained with hematoxylin for 2 min, dehydrated through graded ethanol and xylene, and coverslipped using Cytoseal XYL. Tumor sections were imaged by brightfield microscopy. For YAP and SKP2 stains, rehydration and antigen retrieval were performed the same as 4HNE stains, then permeabilized with 1% Triton-X (Sigma-Aldrich) for 20 min and blocked in 5% BSA for 1hr. Slides were incubated with the YAP or SKP2 antibody (1:200; SantaCruz, #sc-15407 and 1:300 Cell Signaling, #2652 respectively) diluted in appropriate antibody diluent (ThermoFisher Scientific) overnight. After washing with 0.05% Tween-20 in PBS, gels were incubated with Alexa Fluor 546-conjugated secondary antibodies (1:400; Invitrogen, #A11035) for 1 hr, and DAPI (Invitrogen # D1306) for 10 min. Slides were sealed with Fluoromont-G. Tumor sections were imaged by confocal microscopy.

#### Collagen Reflection Microscopy

Reflective microscopy was used to analyze collagen fibers in fresh tumors following embedding in OCT gel and cryosectioning. Images were collected using a Nikon confocal microscope. A 40x objective and 561 nm light were used for reflective microscopy to illuminate and capture collagen fibers.

#### Atomic force microscopy of tumor sections

Fresh non-fixed tumors were resected and embedded in OCT gel and cryosectioned. Before each experiment, the AFM laser was aligned to the back of the cantilever using the Bruker Nanowizard V BioScience optical positioning system to maximize the photodiode sum signal. Alignment was adjusted in both X and Y until the signal reached the manufacturer-recommended range, and the position-sensitive photodetector was centered by adjusting detector offset controls until the vertical and horizontal difference signals were near zero in the free (non-contact) state. Optical lever sensitivity was confirmed during deflection sensitivity calibration on a clean glass substrate. AFM measurements were performed while submerged using a Nanowizard V BioScience (Bruker) equipped with SAA-SPH-1UM probes (spring constant 0.15–0.21 N/m) with a setpoint of 3 nN, velocity of 2 µm/s, and retraction length of 4 µm. Regions for measurement were selected by locating the center of the tumor and moving toward the periphery by approximately 90% of the tumor radius. A 25 × 25 µm scan area was defined, and 16 measurements were acquired within the region at room temperature on a vibration isolation table (TMC CleanBench). The probe oscillated in a frequency shuffle of 1, 2.2, 4.5, 10, 22, 45, and 100 Hz for at least five cycles with an amplitude of 20 nm. The Storage Modulus and Loss Modulus were calculated by fitting the force-displacement curves with an assumed Poisson’s ratio of 0.5 and filtering poorly fitting curves.

### Statistical Analysis

Statistical analyses were performed using GraphPad Prism 10 (GraphPad Software Inc.) or Microsoft Excel, as appropriate. Technical replicates were averaged within each independent biological experiment prior to statistical analysis and were not treated as independent observations. Biological replicates are indicated as N and represent independent experiments performed on separately cultured cell populations, whereas technical replicates are indicated as n and represent repeated measurements within the same biological experiment. Comparisons between two groups were performed using one-way ANOVA. Experiments comparing two conditions across multiple drug concentrations or time points were analyzed using two-way repeated-measures ANOVA. For longitudinal tumor-growth analyses, the same tumor was measured by caliper beginning on the first day of treatment and subsequently every 2–3 days. Tumor volume at each time point was calculated and normalized to the corresponding tumor volume on the first day of treatment, generating a fold change in tumor volume over time to account for differences in initial tumor size. Planned comparisons between treatment groups were analyzed individually using two-way repeated-measures ANOVA to compare longitudinal growth trajectories, with each individual mouse treated as the experimental unit (labeled as N). Multiple-comparisons procedures, if performed, are specified in the corresponding figure legends. All data are presented as mean ± SEM unless otherwise indicated, and P < 0.05 was considered statistically significant.

## Supporting information

All Supplemental Figures

## Data Availability

Data supporting the findings of this study are presented in the paper and the supplementary information. Source data are available through the Duke Research Data Repository (https://research.repository.duke.edu/?ln=en). In accordance with the journal’s reporting requirements, we have included a Supporting Data Values XLS file containing the underlying numerical data for the quantitative analyses presented in the manuscript and supplementary figures.

## Manuscript Information

### Author Contributions

E.V. performed most experiments, designed the study, and wrote the manuscript. N.T. assisted with quantificiation and some experiments. S.W. performed plasmid expansion and isolation of YAPS94A mutant DNA. C.A. performed AFM experiments. J.W. performed SKP2 and YAP western blots. C.L. assisted with mechanistic insight and provided materials necessary for experiments. J.C. and S.G. were the principal investigators of this study and designed the study.

### AI disclosure

OpenAI ChatGPT 5.6 Sol was used between June-October 2026 to assist with language editing, organization, and refinement of manuscript text. AI tools were not used to generate or analyze primary experimental data. The authors reviewed all AI-assisted content and assume full responsibility for the integrity and accuracy of AI output presented in this article.

### Funding Support

This work is the result of National Institutes of Health (NIH) funding, in whole or in part, and is subject to the NIH Public Access Policy. Through acceptance of this federal funding, the NIH has been given a right to make the work publicly available in PubMed Central.

NIH T32 grant 1T32GM144291 (to E.V.)

National Science Foundation GRFP GDS 2139754 (to C.A.)

Duke Cancer Institute (DCI) P30 Cancer Center Support Grant P30CA014236 (to S.G.)

Duke Science and Technology (to S.G.)

DCI Pilot Project (to J.C.)

Paula Takacs Foundation (to S.G. and J.C. jointly)

## Acknowledgements

We thank Emily Warren for the assistance with Western blots. We thank Ssu-Yu Chen for training in the necessary assays. We thank YingYu Lin and Jiyeon Song for their guidance with animal work.

## Address for correspondence

Jen-Tsan Ashley Chi, Duke Box 3382, Durham, NC 27710

Sharon Gerecht, Duke Box 90281, Durham, NC 27708

---

**Supplemental Figure 1:**
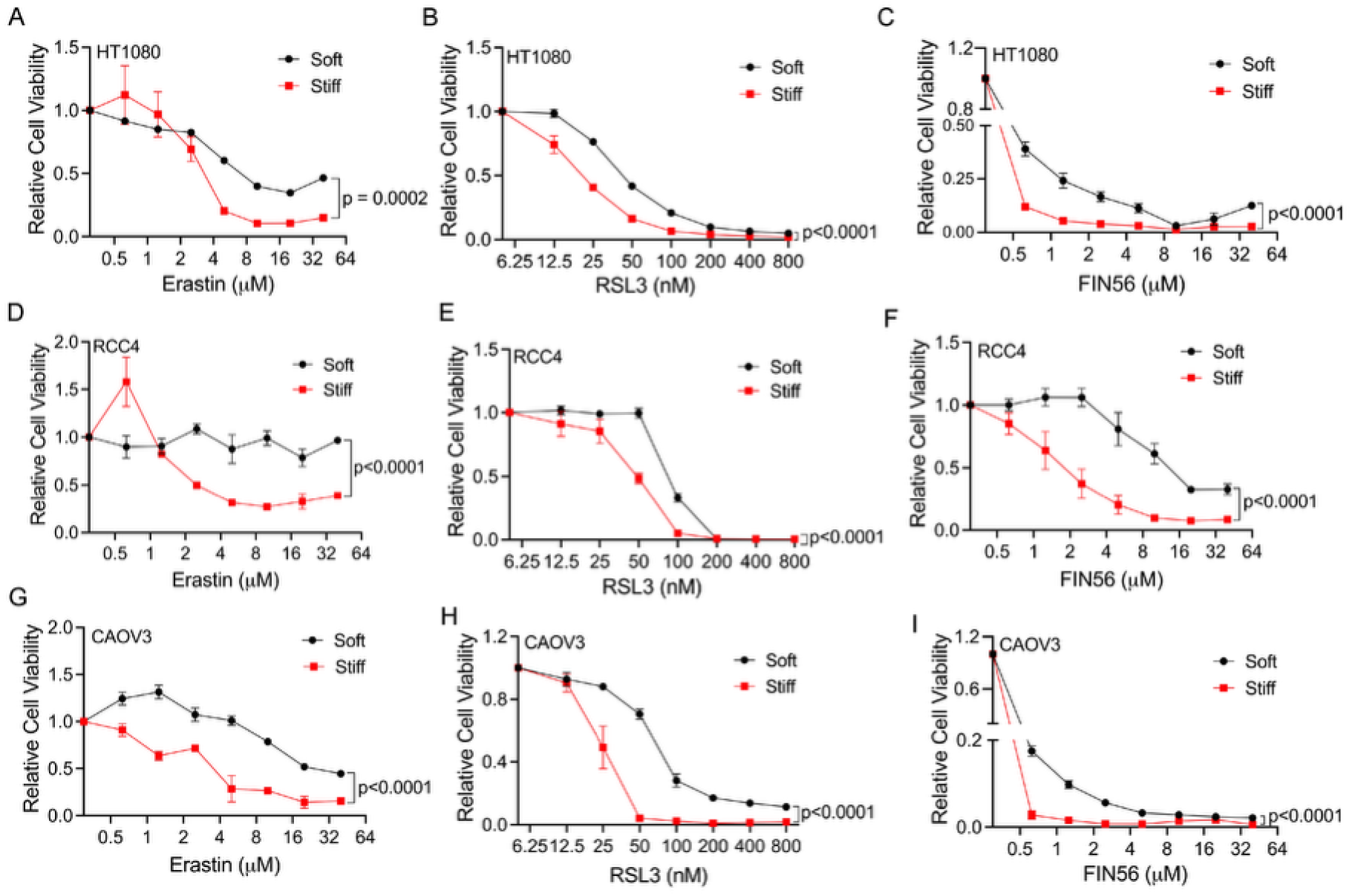
Ferroptosis is regulated by matrix stiffness in multiple cancer types. (**A**) Relative cell viability of HT1080 fibrosarcoma cancer cells on soft or stiff hydrogels treated with erastin, (**B**) RSL3, and (**C**) FIN56 for 16 hr. (N=4) (**D**) Relative cell viability of RCC4 renal cancer cells on soft or stiff hydrogels treated with erastin, (**E**) RSL3, or (**F**) FIN56 for 16 hr. (N=3) (**G**) Relative cell viability of CAOV3 ovarian cancer cells on soft or stiff hydrogels treated with erastin, (**H)** FIN56, or (**I**) RSL3 for 16 hr (N=3) All graphs represent means ± SEM. Two-way repeated-measures ANOVA was used for A-I.

**Supplemental Figure 2:**
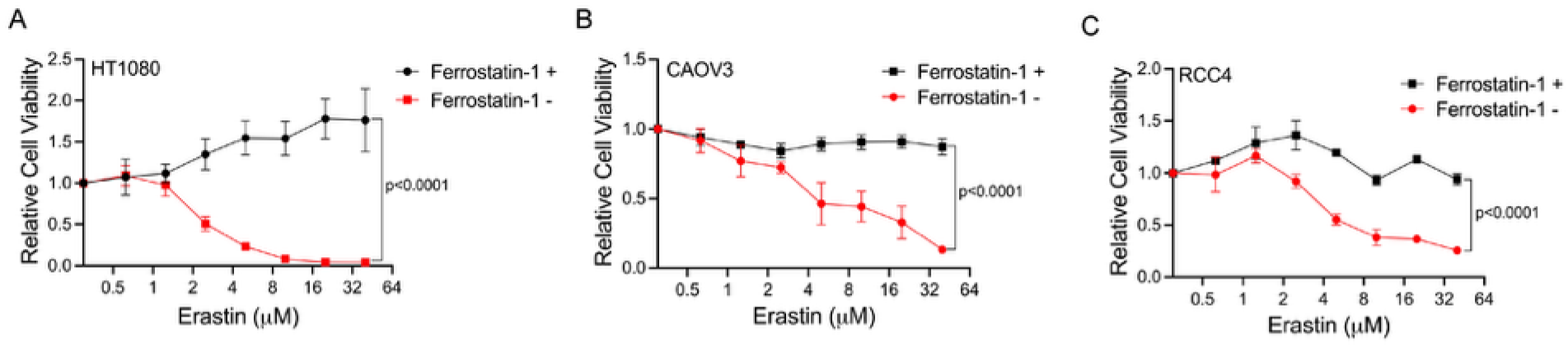
Ferrostatin-1 rescues erastin-induced ferroptotic death. (**A**) HT1080 (**B**) CAOV3, or (**C**) RCC4 cells on stiff hydrogels treated with erastin with or without 6 μM ferrostatin-1 (ferroptosis inhibitor) for 16 hr. (N=3) All graphs represent means ± SEM. Two-way repeated-measures ANOVA was used for A-C.

**Supplemental Figure 3:**
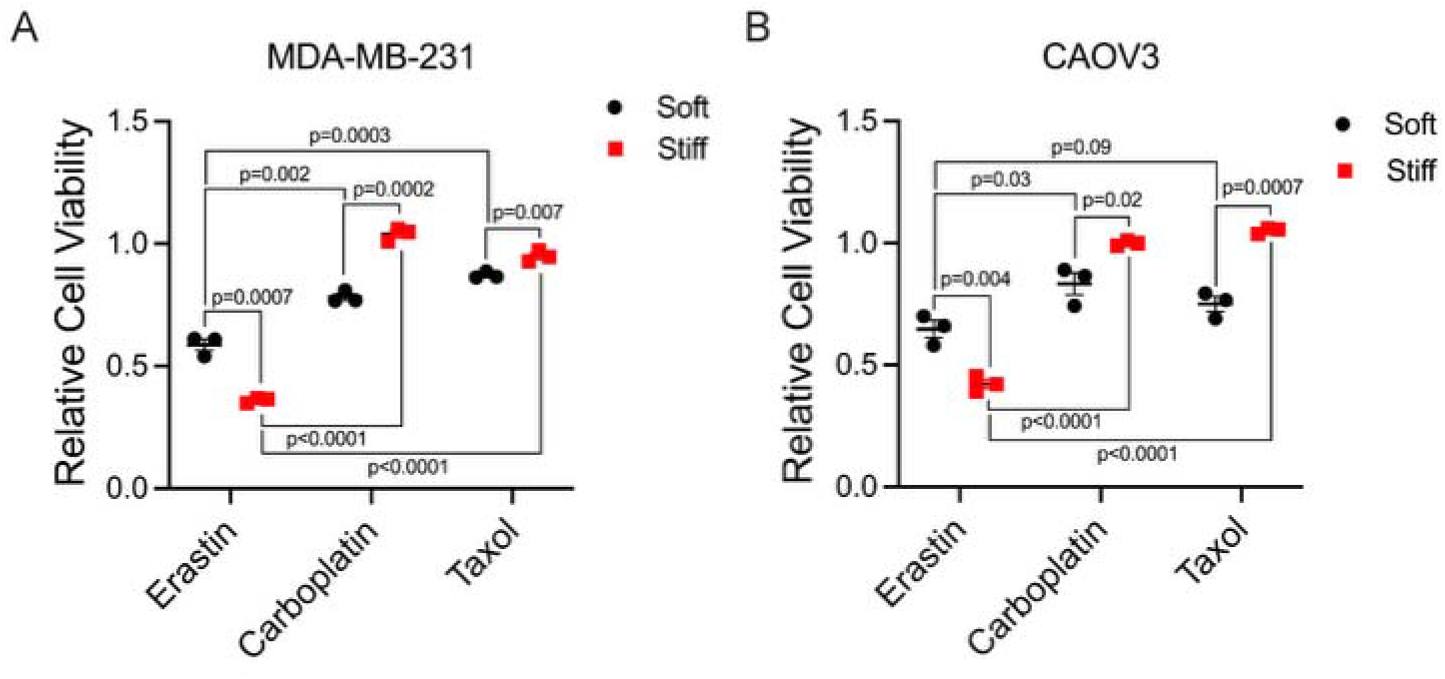
MDA-MB-231 and CAOV3 cells seeded on stiff hydrogels are chemoresistant. (**A**) Relative cell viability of MDA-MB-231 or (**B**) CAOV3 cells seeded on stiff or soft hydrogels treated for 24 hr with 200 μM carboplatin, 10nM Taxol, or 10 μM erastin (N=3). Lines in graphs represent mean. One-way ANOVA was measured separately for each comparison in A and B.

**Supplemental Figure 4:**
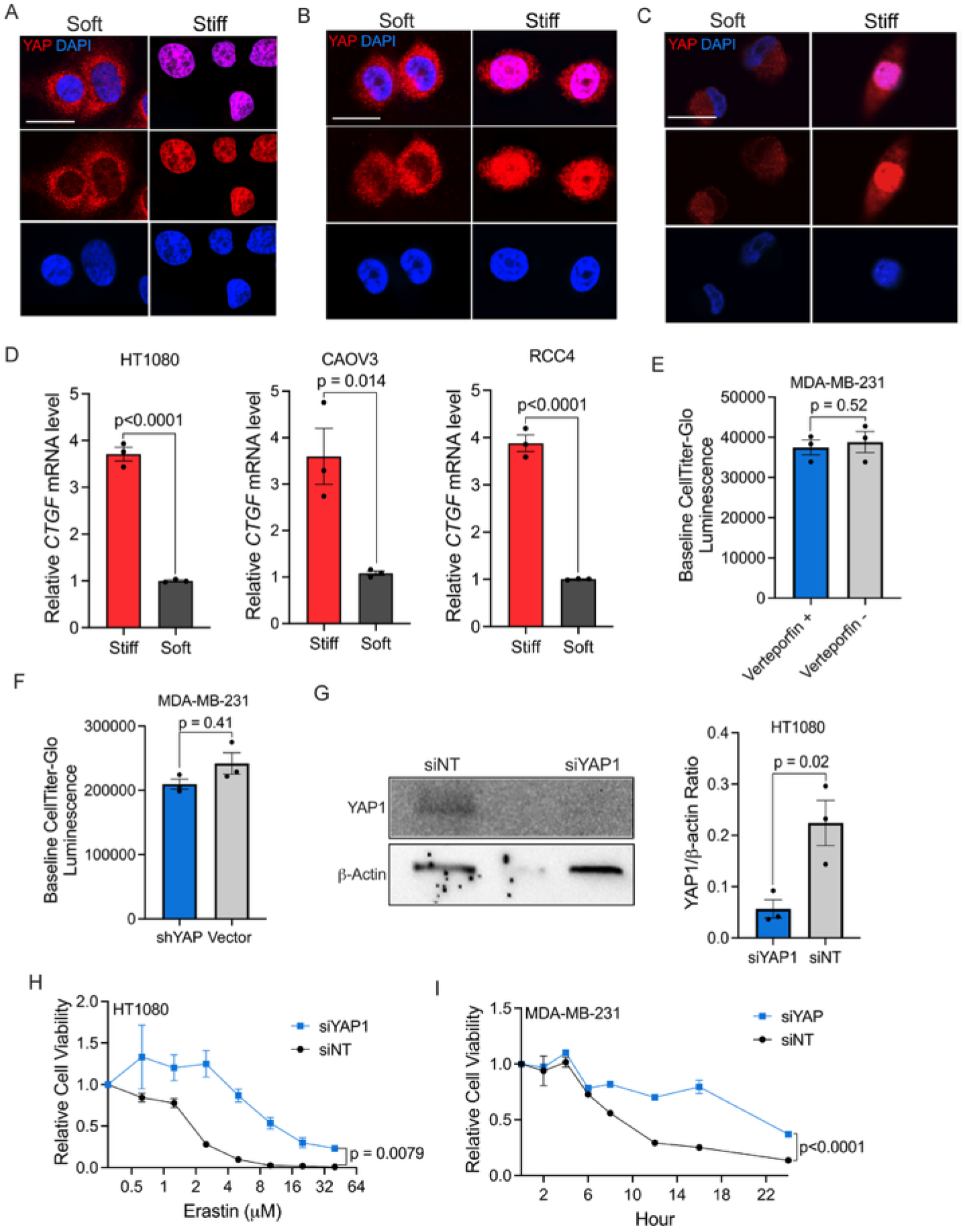
Matrix-mediated ferroptosis is regulated by YAP. (**A**) Representative immunofluorescence images of HT1080 cells, (**B**) CAOV3 cells, or (**C**) RCC4 cells stained with DAPI (nuclei in blue) and YAP (in red). Scale bars=25 μm (Representative images of N=3 independent experiments) (**D**) Relative mRNA fold change of *CTGF* measured by qRT-PCR of HT1080, CAOV3 and RCC4 cells. (**E**) Baseline CellTtiter-Glo luminescence reading of 8 μM verteporfin and DMSO controls after 16 hr treatment. (N=3) (**F**) Baseline CellTtiter-Glo luminescence reading of shYAP and vector controls with no treatment. (N=3) (**G**) Western blot of siYAP1 and siNT transfected HT1080 cells stained for YAP and B-Actin (left) and quantification (right). (N=3) (**H**) Relative cell viability of siYAP1 or siNT HT1080 cells treated with erastin for 16 hr. (N=3) (**I**) Relative cell viability of MDA-MB-231 cells treated with either siYAP or siNT then treated with 10 μM erastin and measured for viability at 0, 2, 4, 6, 8, 12, 16 and 24hr on stiff hydrogels. (N=3) All graphs represent means ± SEM. One-way ANOVA was used for D-G. Two-way repeated-measures ANOVA was used for H and I.

**Supplemental Figure 5:**
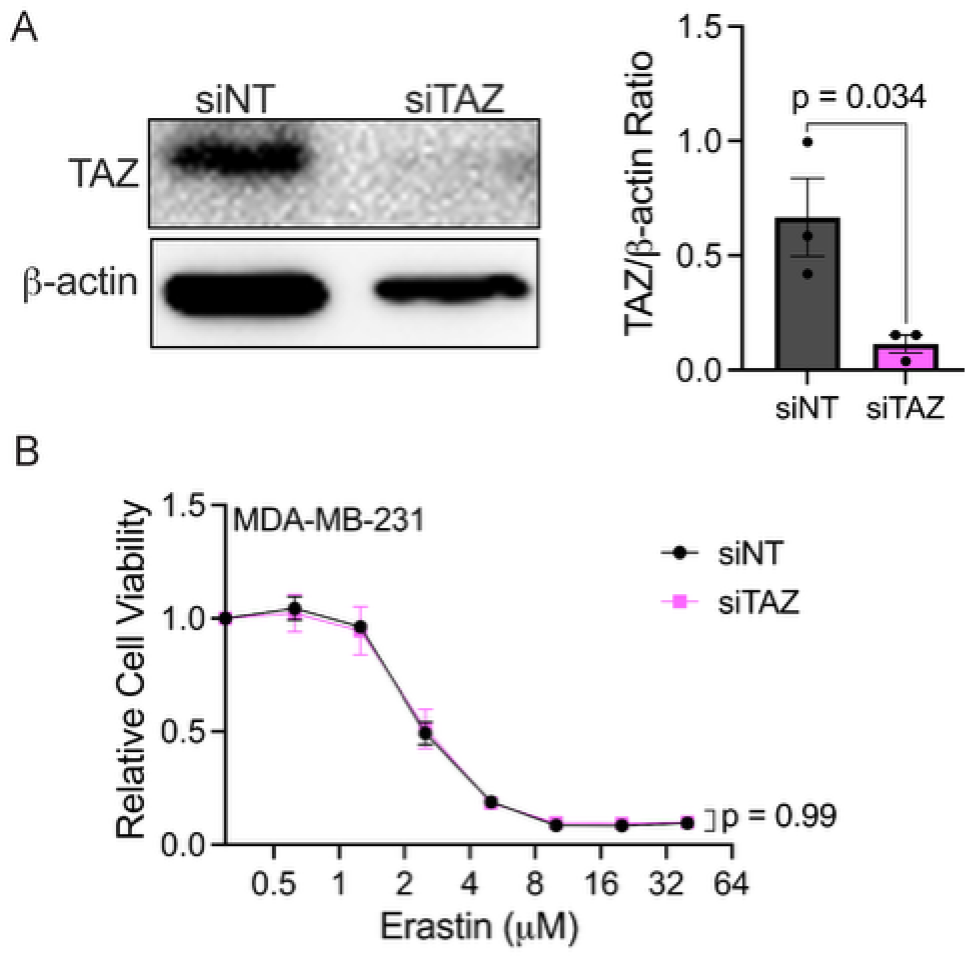
Matrix-mediated ferroptosis is not regulated by TAZ. (**A**) Western blot of siTAZ and siNT transfected MDA-MB-231 cells stained for TAZ and B-Actin (left) and quantification (right). (N=3) (**B**) Relative cell viability of MDA-MB-231 cells transfected with either siNT or siTAZ treated with erastin for 16 hr on stiff hydrogels. (N=3). All graphs represent means ± SEM. One-way ANOVA was used for A. Two-way repeated-measures ANOVA was used for B.

**Supplemental Figure 6:**
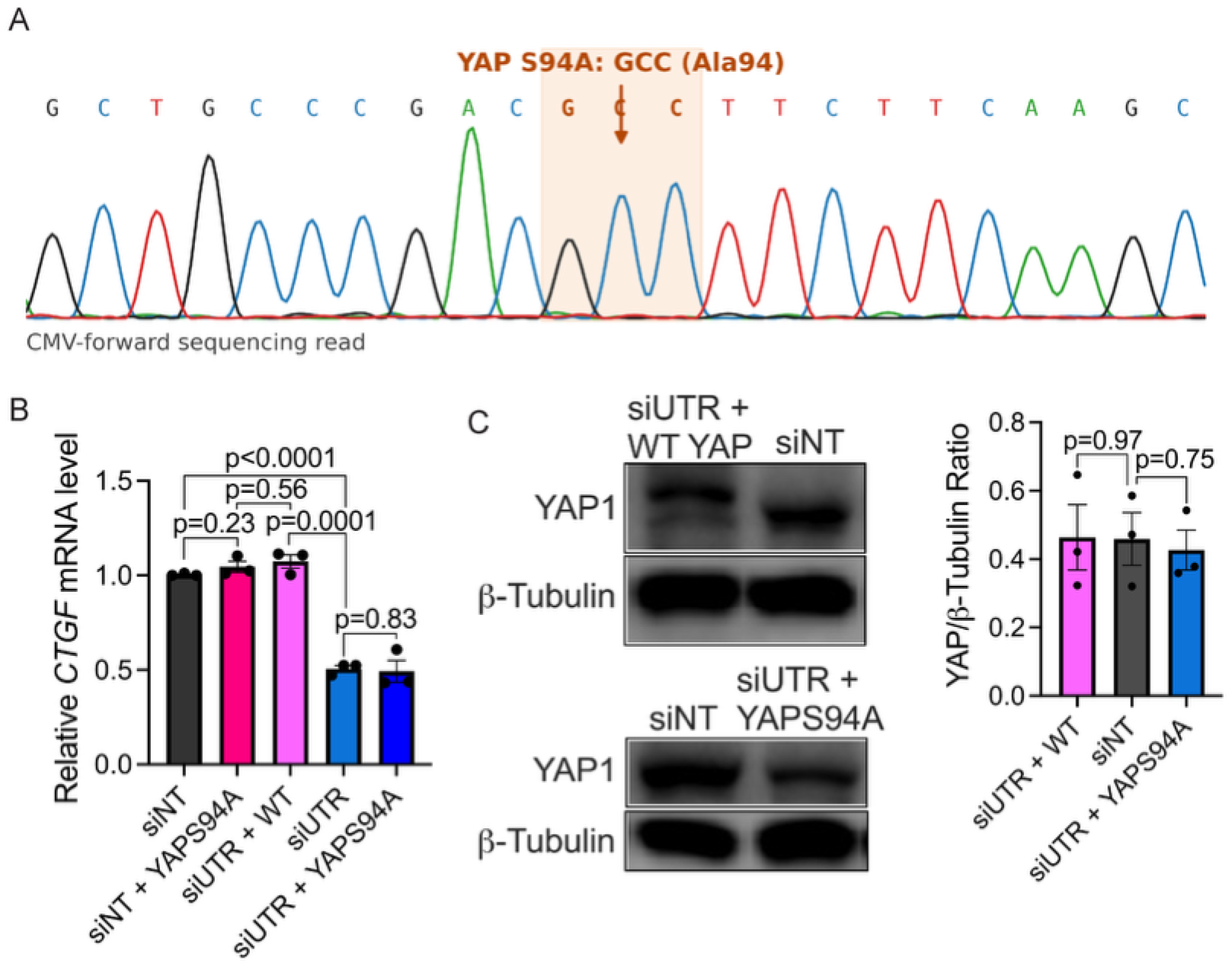
YAPS94A mutant confirmation. (**A**) Sanger sequencing confirming YAPS94A mutation. (**B**) Relative *CTGF* mRNA level measured by RT-PCR for each condition. (N=3) (**C**) Western blot of siNT, siUTR+YAPS94A, and siUTR+WT YAP transfected MDA-MB-231 cells stained for YAP and GAPDH (left) and quantification (right). (N=3) All bar graphs represent ± SEM. One-way ANOVA was used for B and C.

**Supplemental Figure 7:**
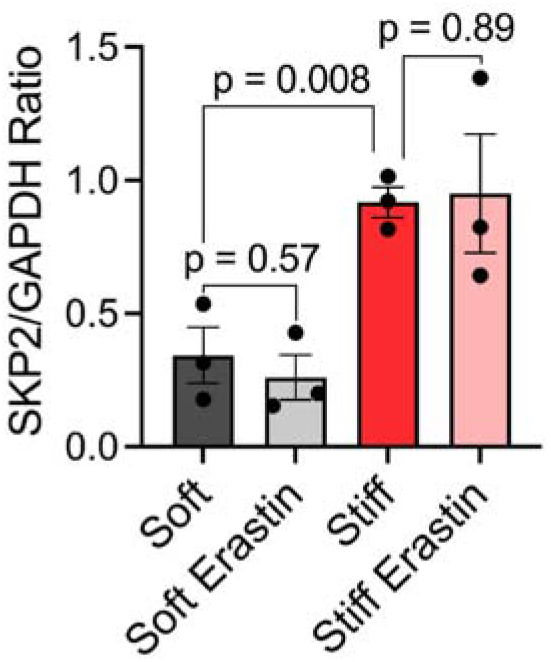
SKP2 western blot quantification. Quantification of western blots of MDA-MB-231 cells seeded on stiff or soft hydrogels treated with DMSO or erastin for 12 hr stained for SKP2 or GAPDH (N=3) Bar graph represents means ± SEM. One-way ANOVA was used.

**Supplemental Figure 8:**
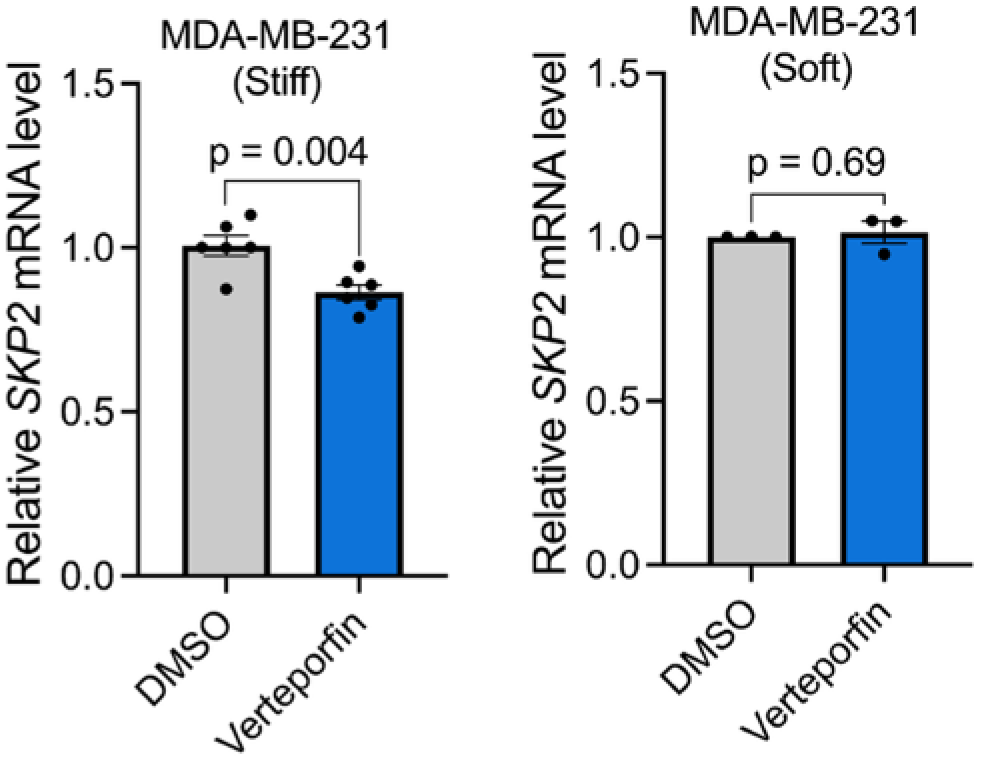
*SKP2* mRNA fold change after verteporfin. Relative *SKP2* mRNA level of MDA-MB-231 cells seeded on soft or stiff hydrogels treated with DMSO or 8 μM verteporfin for 24 hr. (left- N=6 and right N=3). Bar graph represents means ± SEM. One-way ANOVA was used.

**Supplemental Figure 9:**
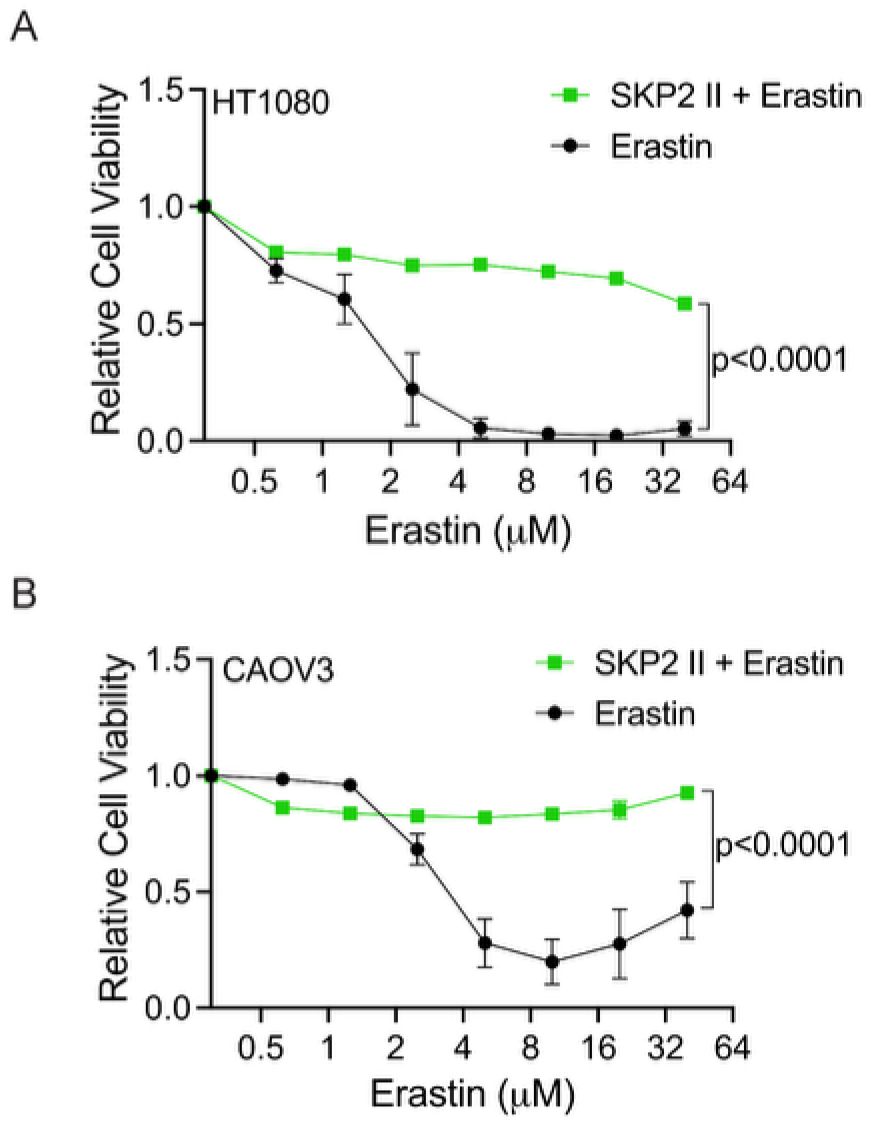
SKP2 inhibition rescues cell viability in other cell lines. (**A**) Relative cell viability with erastin only or combined with 40 μM SKP2 inhibition in HT1080 cells or (**B**) CAOV3 cells for 16 hr. (N=3 for all panels) All graphs represent means ± SEM. Two-way repeated-measures ANOVA was used.

**Supplemental Figure 10:**
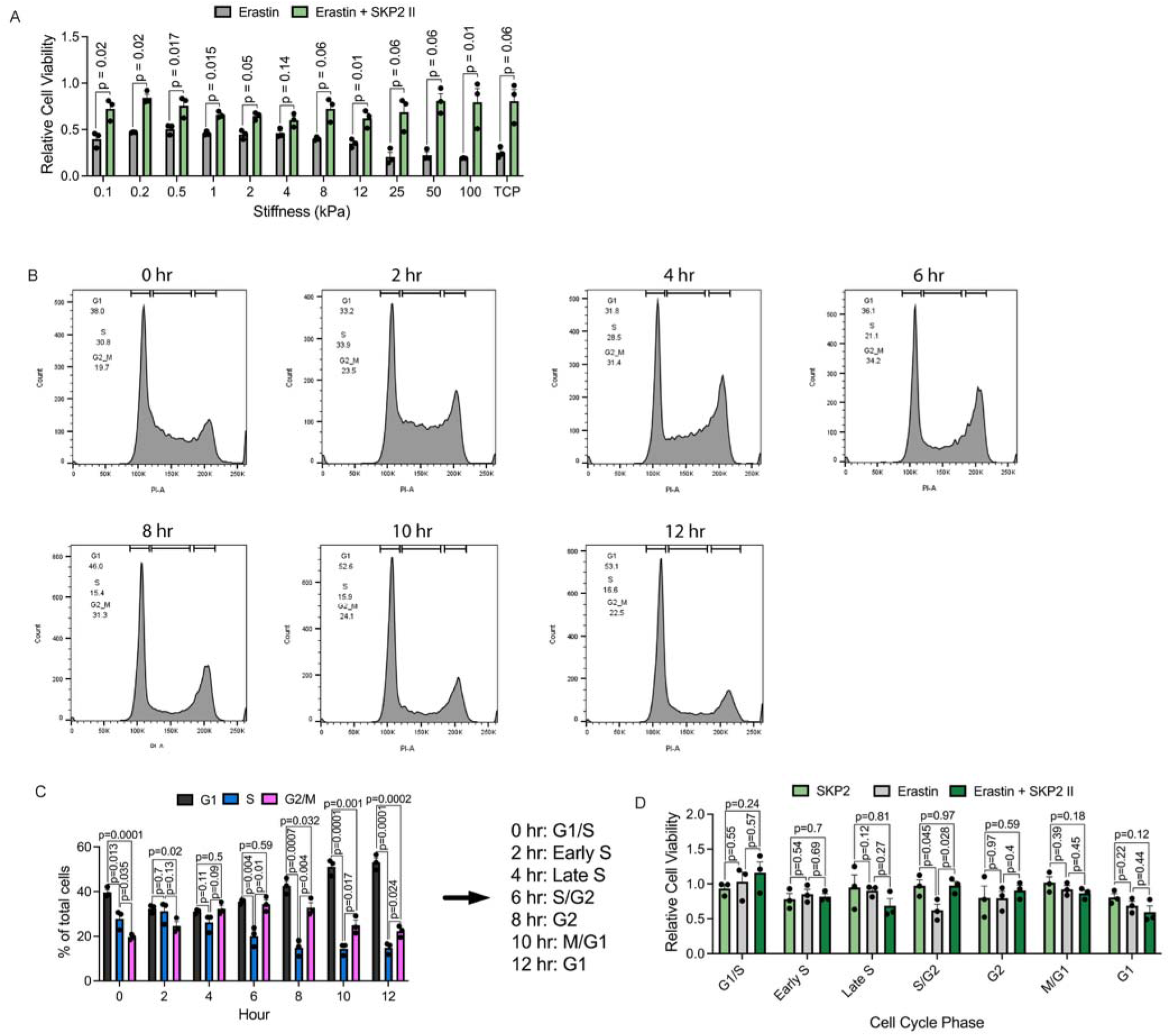
Effect of stiffness and cell cycle on SKP2-mediated ferroptosis. (**A**) Relative cell viability of MDA-MB-231 cells seeded on hydrogels of various stiffnesses or tissue culture plastic (TCP) treated with 10 μM erastin alone or erastin and 40 μM SKP2 inhibitor II for 16 hr. (N=3) (**B**) Representative flow cytometry histograms of PI flow every 2 hr after cell cycle synchronization with gating. (N=3) (**C**) Quantification of PI flow for G1, S, and G2/M phases for cell synchronization and labels given for each time point. (N=3) (**D**) Relative cell viability of MDA-MB-231 cells for each cell cycle phase following treatment with 40 μM SKP2 inhibitor II, 10 μM erastin, or combination for 6 hr. (N=3) All graphs represent means ± SEM. One-way ANOVA was used for each comparison.

**Supplemental Figure 11:**
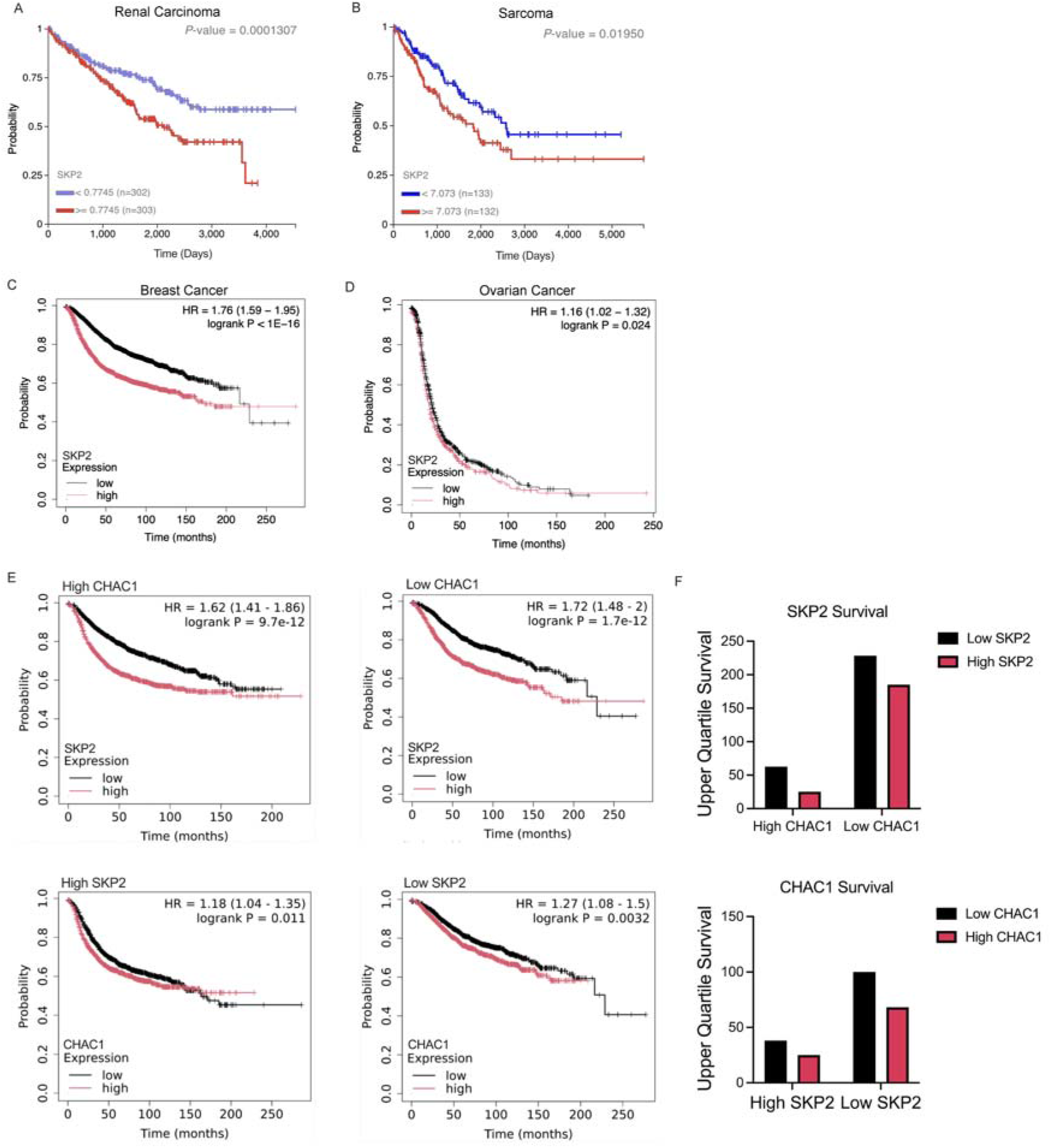
Survival data for *SKP2* high and low tumors. (**A**) Kaplan Meier plots for overall survival by *SKP2* low or high expression in renal carcinoma (45) (Low: N=302; High: N=303) (**B**) overall survival in sarcoma (45) (Low: N=133; High: N=132), (**C**) recurrence-free survival in breast cancer (44) (Low: N=2462; High: N=2467), (**D**) progression-free survival ovarian cancer (44) (Low: N=714; High: N=715). (**E**) Kaplan Meier plots for recurrence-free survival in breast cancer by *SKP2* low or high expression or *CHAC1* low or high expression filtered by low or high *SKP2* or *CHAC1* expression (44). (High *CHAC1*, Low *SKP2*: N=1220 High *SKP2*: N=1223; Low *CHAC1*, Low *SKP2*: N=1228 High *SKP2*: N=1221; High *SKP2*, Low *CHAC1*: N=1235, High *CHAC1*: N=1227; Low *SKP2*, Low *CHAC1*: N=1239 High *CHAC1*: N=1221), (**F**) Upper quartile survival quantification of Kaplan Meier plots for survival.

**Supplemental Figure 12:**
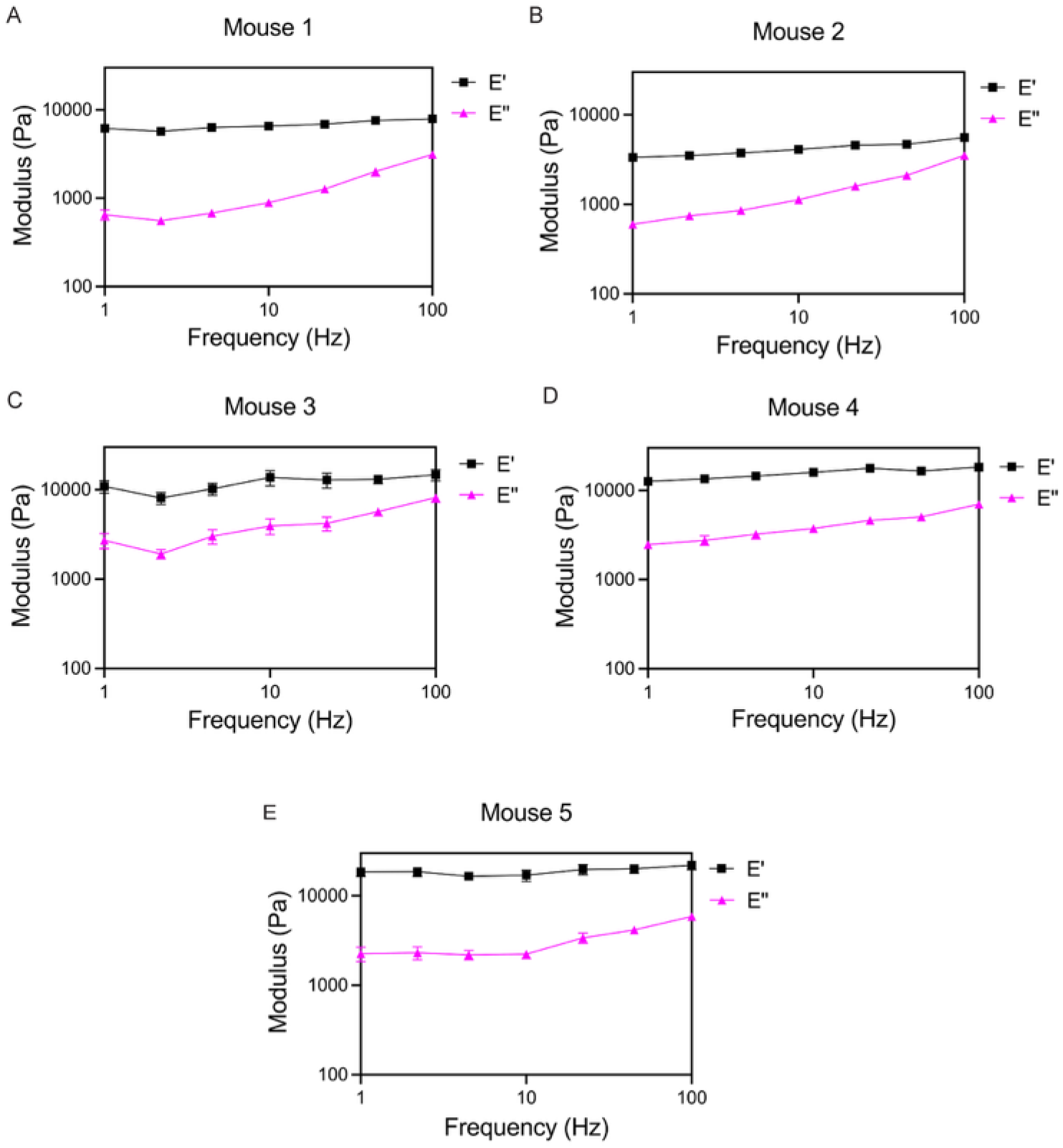
AFM frequency sweep of individual tumors. (A-E) Frequency sweep for storage modulus E’ and loss modulus E’’ measured by AFM for PBS treated mice 1-5.

**Supplemental Figure 13:**
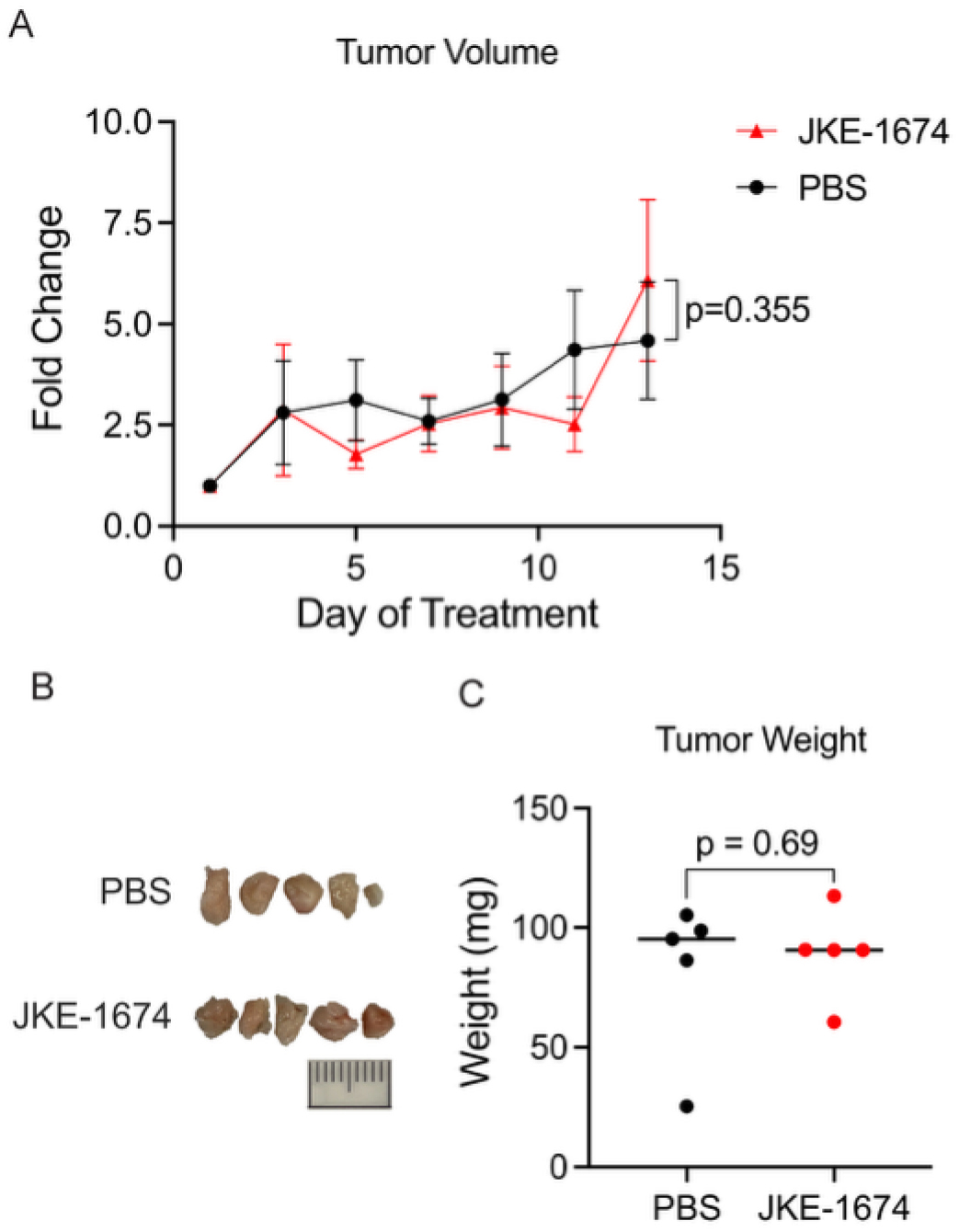
Ferroptosis lacks therapeutic potential for soft breast tumors. (A) Fold change in tumor volume, measured by calipers, relative to the initial tumor volume on day 7 after treatment initiation. (B) Images of isolated tumors for each condition. Scale bar = 10 mm (C) Measured weight of whole isolated tumors for each condition. (N=5 for all conditions and measurements). Figure 1A graph represents means ± SEM. Lines in graph in C represent mean. One-way ANOVA was used for C. Two-way repeated-measures ANOVA was used for A.

**Supplemental Figure 14:**
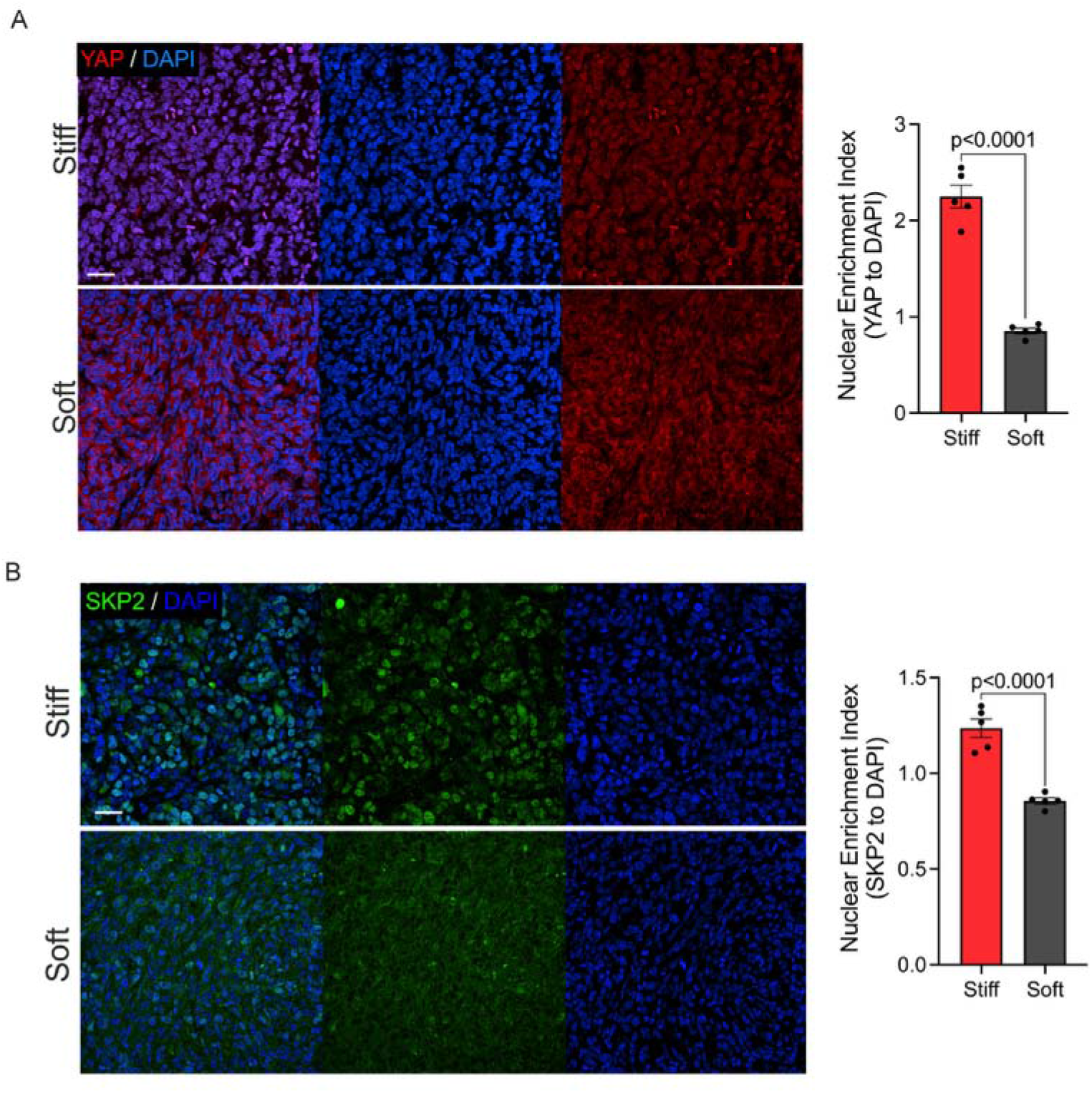
YAP and SKP2 stains for soft and stiffened tumors. (A) Representative immunofluorescence images of MDA-MB-231 soft or stiff generated tumors stained with DAPI (nuclei in blue) and YAP (in red) (left) and quantification (right). Scale bar=25 μm. Representative image shown; analysis was performed on tumors from N = 5 mice per group. (B) Representative immunofluorescence images of MDA-MB-231 soft or stiff generated tumors stained with DAPI (nuclei in blue) and SKP2 (in green) (left) and quantification (right). Scale bar=25 μm. Representative image shown; analysis was performed on tumors from N = 5 mice per group. Nuclear enrichment index >1 indicates nuclear enrichment and <1 nuclear exclusion. All graphs represent means ± SEM. One-way ANOVA was used for analysis.

**Supplemental Figure 15:**
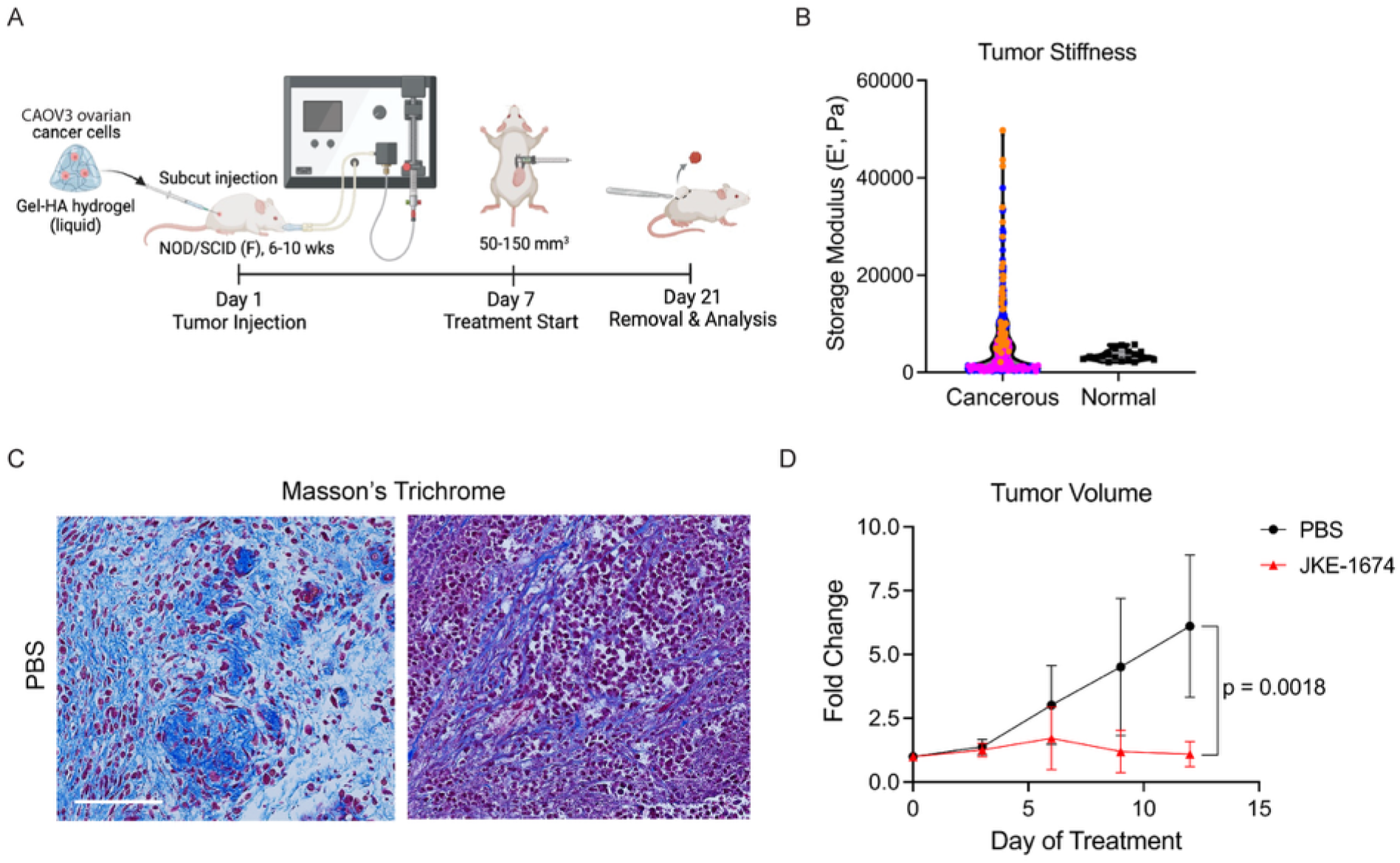
Ferroptosis can target stiff ovarian tumors. (A) Schematic of experimental timeline for establishing stiff ovarian tumors in vivo. (B) AFM measurements of fresh ovarian tumor sections compared to normal ovarian tissue from (52). (C) Representative images of Masson’s trichrome staining of PBS treated tumor sections. Scale bars=100 μm (D) Fold change in tumor volume, measured by calipers, relative to the initial tumor volume on day 7 after treatment initiation. (N=3) All graphs represent means ± SEM except B. Two-way repeated-measures ANOVA was used for D.

