## Supplementary material for "Tumor stiffness creates a ferroptotic Achilles’ heel through YAP–SKP2 signaling": All Supplemental Figures

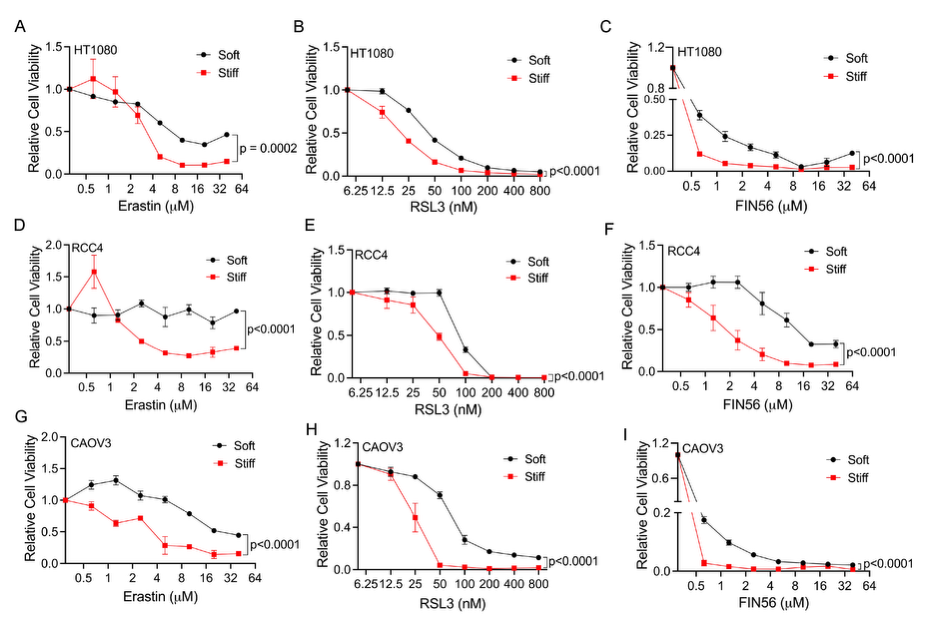

**Supplemental Figure 1: Ferroptosis is regulated by matrix stiffness in multiple cancer types.** (**A**) Relative cell viability of HT1080 fibrosarcoma cancer cells on soft or stiff hydrogels treated with erastin**,** (**B**) RSL3, and (**C**) FIN56 for 16 hr. (N=4) (**D**) Relative cell viability of RCC4 renal cancer cells on soft or stiff hydrogels treated with erastin, (**E**) RSL3, or (**F**) FIN56 for 16 hr. (N=3) (**G**) Relative cell viability of CAOV3 ovarian cancer cells on soft or stiff hydrogels treated with erastin, (**H)** FIN56, or (**I**) RSL3 for 16 hr (N=3) All graphs represent means ± SEM. Two-way repeated-measures ANOVA was used for A-I.

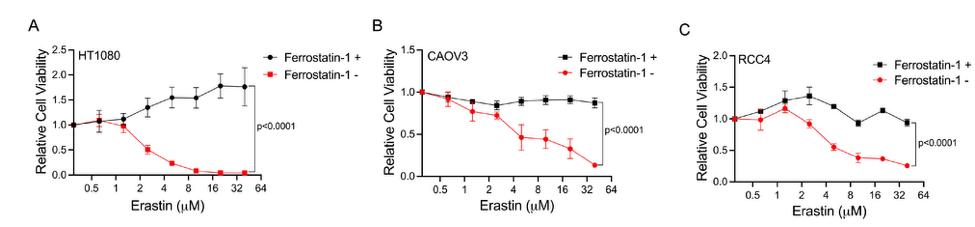

**Supplemental Figure 2:** **Ferrostatin-1 rescues erastin-induced ferroptotic death.** (**A**) HT1080 (**B**) CAOV3, or (**C**) RCC4 cells on stiff hydrogels treated with erastin with or without 6 $\mu$M ferrostatin-1 (ferroptosis inhibitor) for 16 hr. (N=3) All graphs represent means ± SEM. Two-way repeated-measures ANOVA was used for A-C.

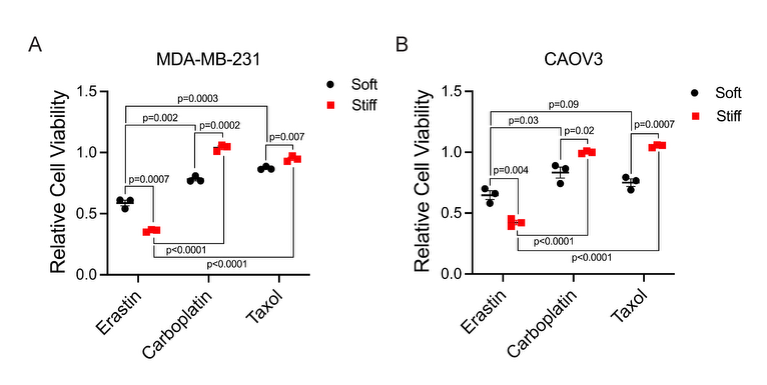

**Supplemental Figure 3: MDA-MB-231 and CAOV3 cells seeded on stiff hydrogels are chemoresistant.** (**A**) Relative cell viability of MDA-MB-231 or (**B**) CAOV3 cells seeded on stiff or soft hydrogels treated for 24 hr with 200 $\mu$M carboplatin, 10nM Taxol, or 10 $\mu$M erastin (N=3). Lines in graphs represent mean. One-way ANOVA was measured separately for each comparison in A and B.

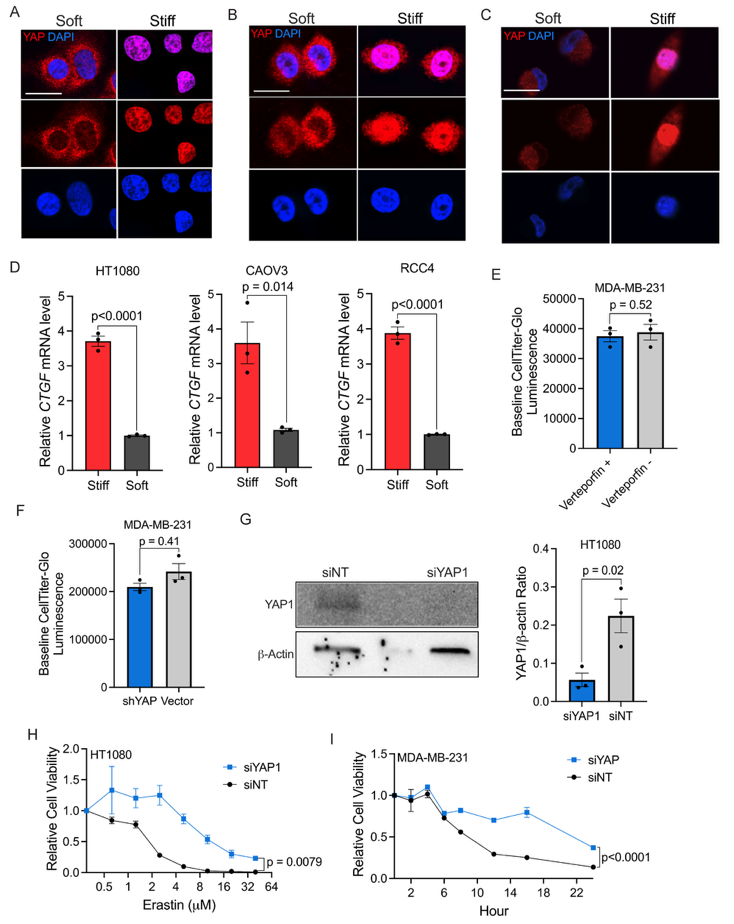

**Supplemental Figure 4: Matrix-mediated ferroptosis is regulated by YAP.** (**A**) Representative immunofluorescence images of HT1080 cells, (**B**) CAOV3 cells, or (**C**) RCC4 cells stained with DAPI (nuclei in blue) and YAP (in red). Scale bars=25 $\mu m$ (Representative images of N=3 independent experiments) (**D**) Relative mRNA fold change of *CTGF* measured by qRT-PCR of HT1080, CAOV3 and RCC4 cells. (**E**) Baseline CellTtiter-Glo luminescence reading of 8 $\mu$M verteporfin and DMSO controls after 16 hr treatment. (N=3) (**F**) Baseline CellTtiter-Glo luminescence reading of shYAP and vector controls with no treatment. (N=3) (**G**) Western blot of siYAP1 and siNT transfected HT1080 cells stained for YAP and B-Actin (left) and quantification (right). (N=3) (**H**) Relative cell viability of siYAP1 or siNT HT1080 cells treated with erastin for 16 hr. (N=3) (**I**) Relative cell viability of MDA-MB-231 cells treated with either siYAP or siNT then treated with 10 $\mu$M erastin and measured for viability at 0, 2, 4, 6, 8, 12, 16 and 24hr on stiff hydrogels. (N=3) All graphs represent means ± SEM. One-way ANOVA was used for D-G. Two-way repeated-measures ANOVA was used for H and I.

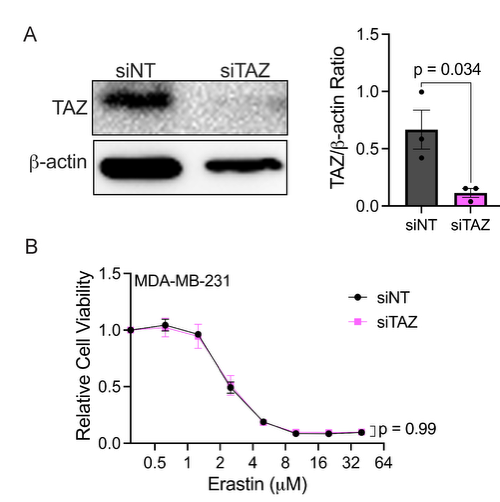

**Supplemental Figure 5: Matrix-mediated ferroptosis is not regulated by TAZ.** (**A**) Western blot of siTAZ and siNT transfected MDA-MB-231 cells stained for TAZ and B-Actin (left) and quantification (right). (N=3) (**B**) Relative cell viability of MDA-MB-231 cells transfected with either siNT or siTAZ treated with erastin for 16 hr on stiff hydrogels. (N=3). All graphs represent means ± SEM. One-way ANOVA was used for A. Two-way repeated-measures ANOVA was used for B.

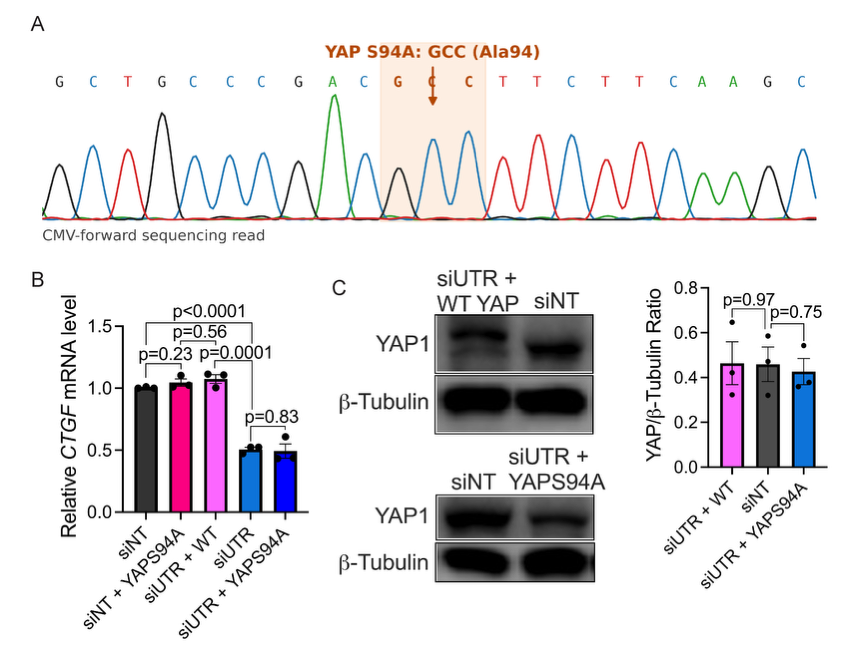

**Supplemental Figure 6: YAPS94A mutant confirmation.** (**A**) Sanger sequencing confirming YAPS94A mutation. (**B**) Relative *CTGF* mRNA level measured by RT-PCR for each condition. (N=3) (**C**) Western blot of siNT, siUTR+YAPS94A, and siUTR+WT YAP transfected MDA-MB-231 cells stained for YAP and GAPDH (left) and quantification (right). (N=3) All bar graphs represent ± SEM. One-way ANOVA was used for B and C.

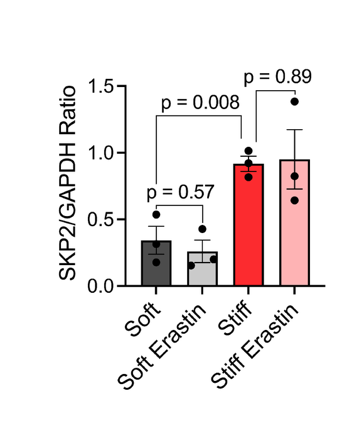

**Supplemental Figure 7: SKP2 western blot quantification.** Quantification of western blots of MDA-MB-231 cells seeded on stiff or soft hydrogels treated with DMSO or erastin for 12 hr stained for SKP2 or GAPDH (N=3) Bar graph represents means ± SEM. One-way ANOVA was used.

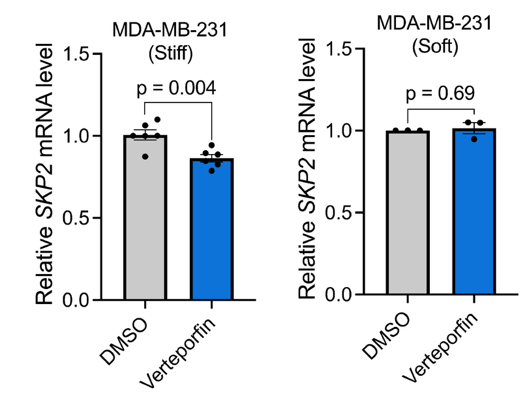

**Supplemental Figure 8: *SKP2* mRNA fold change after verteporfin.** Relative *SKP2* mRNA level of MDA-MB-231 cells seeded on soft or stiff hydrogels treated with DMSO or 8 $\mu$M verteporfin for 24 hr. (left- N=6 and right N=3). Bar graph represents means ± SEM. One-way ANOVA was used.

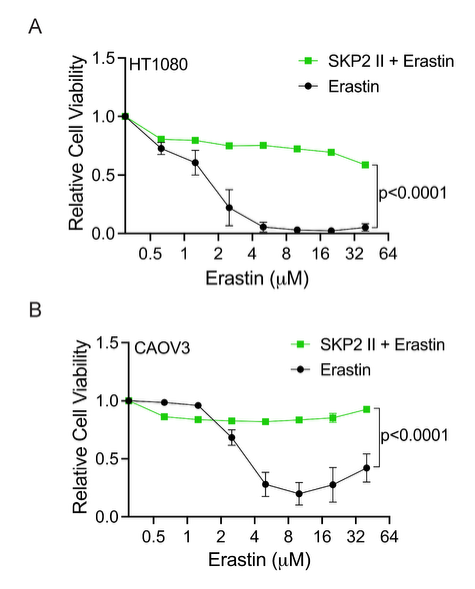

**Supplemental Figure 9: SKP2 inhibition rescues cell viability in other cell lines.** (**A**) Relative cell viability with erastin only or combined with 40 $\mu M$ SKP2 inhibition in HT1080 cells or (**B**) CAOV3 cells for 16 hr. (N=3 for all panels) All graphs represent means ± SEM. Two-way repeated-measures ANOVA was used.

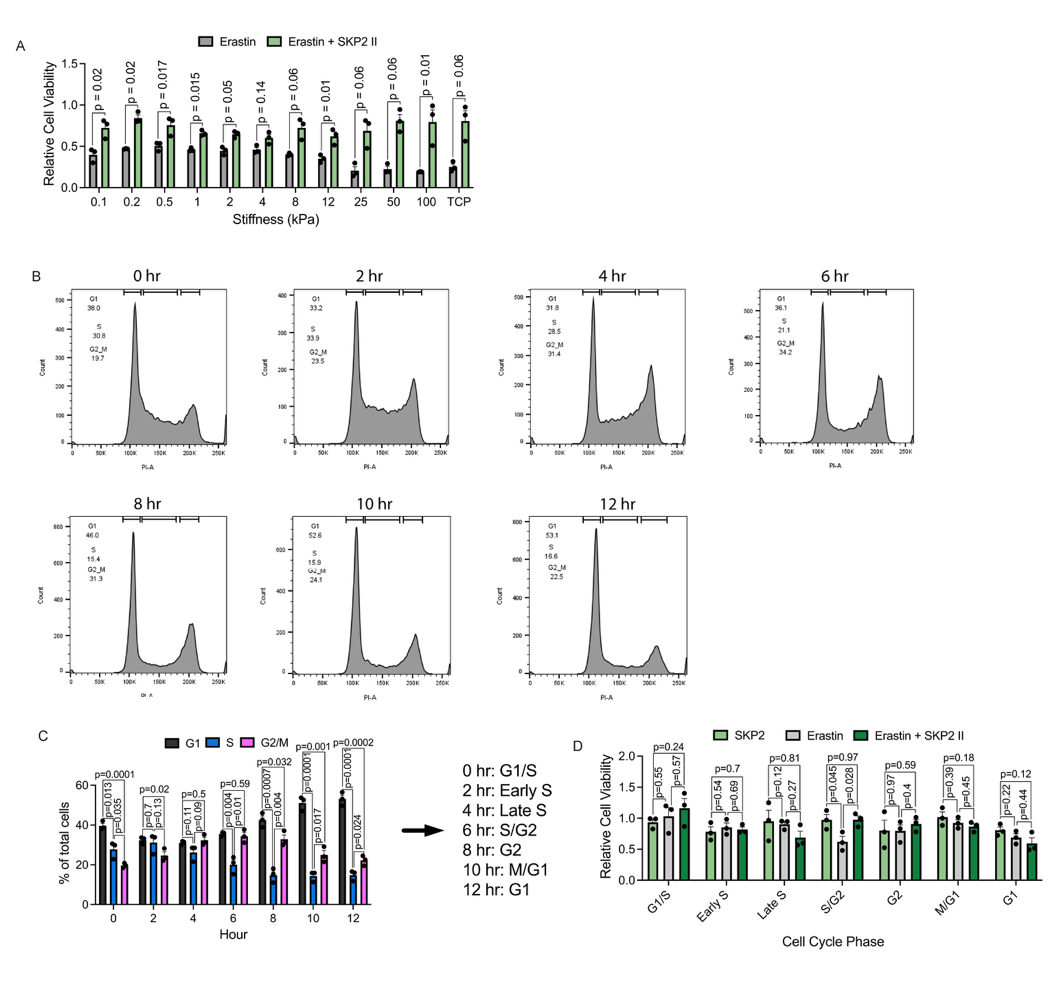
**Supplemental Figure 10: Effect of stiffness and cell cycle on SKP2-mediated ferroptosis.** (**A**) Relative cell viability of MDA-MB-231 cells seeded on hydrogels of various stiffnesses or tissue culture plastic (TCP) treated with 10 $\mu M$erastin alone or erastin and 40 $\mu M$ SKP2 inhibitor II for 16 hr. (N=3) (**B**) Representative flow cytometry histograms of PI flow every 2 hr after cell cycle synchronization with gating. (N=3) (**C**) Quantification of PI flow for G1, S, and G2/M phases for cell synchronization and labels given for each time point. (N=3) (**D**) Relative cell viability of MDA-MB-231 cells for each cell cycle phase following treatment with 40 $\mu M$SKP2 inhibitor II, 10 $\mu M$erastin, or combination for 6 hr. (N=3) All graphs represent means ± SEM. One-way ANOVA was used for each comparison.

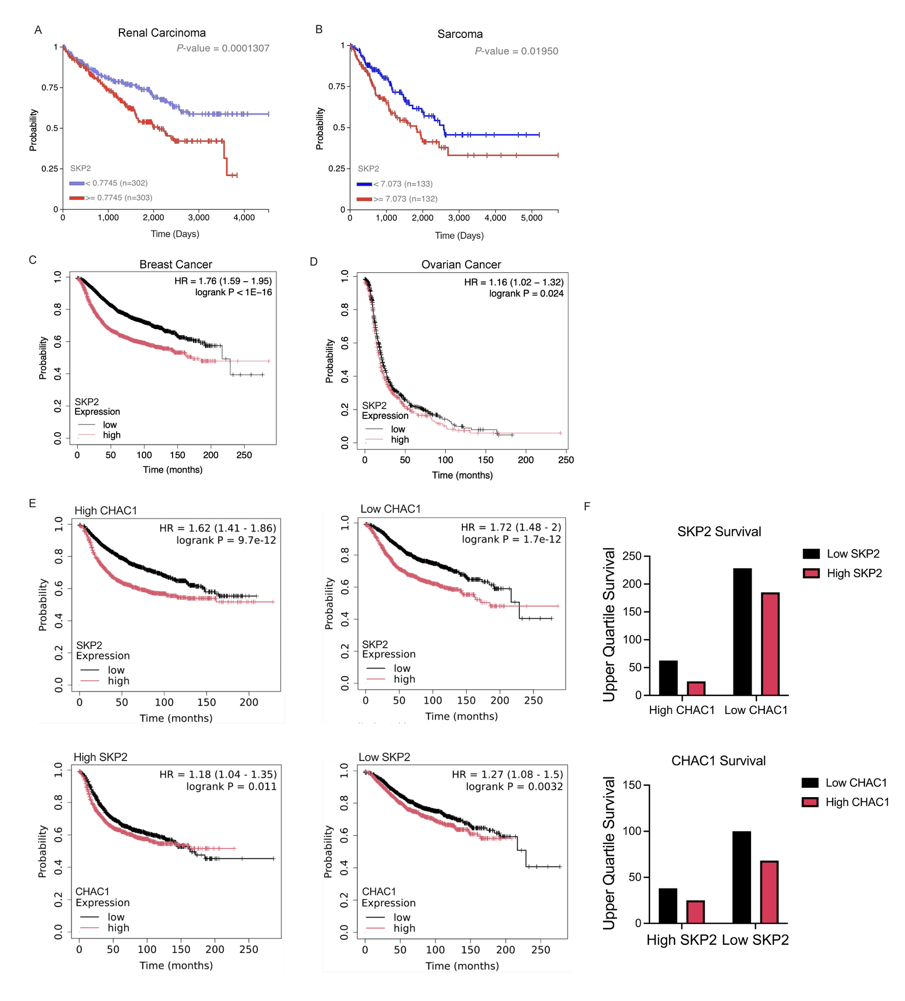
**Supplemental Figure 11: Survival data for *SKP2* high and low tumors.** (**A**) Kaplan Meier plots for overall survival by *SKP2* low or high expression in renal carcinoma (45) (Low: N=302; High: N=303) (**B**) overall survival in sarcoma (45) (Low: N=133; High: N=132), (**C**) recurrence-free survival in breast cancer (44) (Low: N=2462; High: N=2467), (**D**) progression-free survival ovarian cancer (44) (Low: N=714; High: N=715). (**E**) Kaplan Meier plots for recurrence-free survival in breast cancer by *SKP2* low or high expression or *CHAC1* low or high expression filtered by low or high *SKP2* or *CHAC1* expression (44). (High *CHAC1*, Low *SKP2*: N=1220 High *SKP2*: N=1223; Low *CHAC1*, Low *SKP2*: N=1228 High *SKP2*: N=1221; High *SKP2*, Low *CHAC1*: N=1235, High *CHAC1*: N=1227; Low *SKP2*, Low *CHAC1*: N=1239 High *CHAC1*: N=1221), (**F**) Upper quartile survival quantification of Kaplan Meier plots for survival.

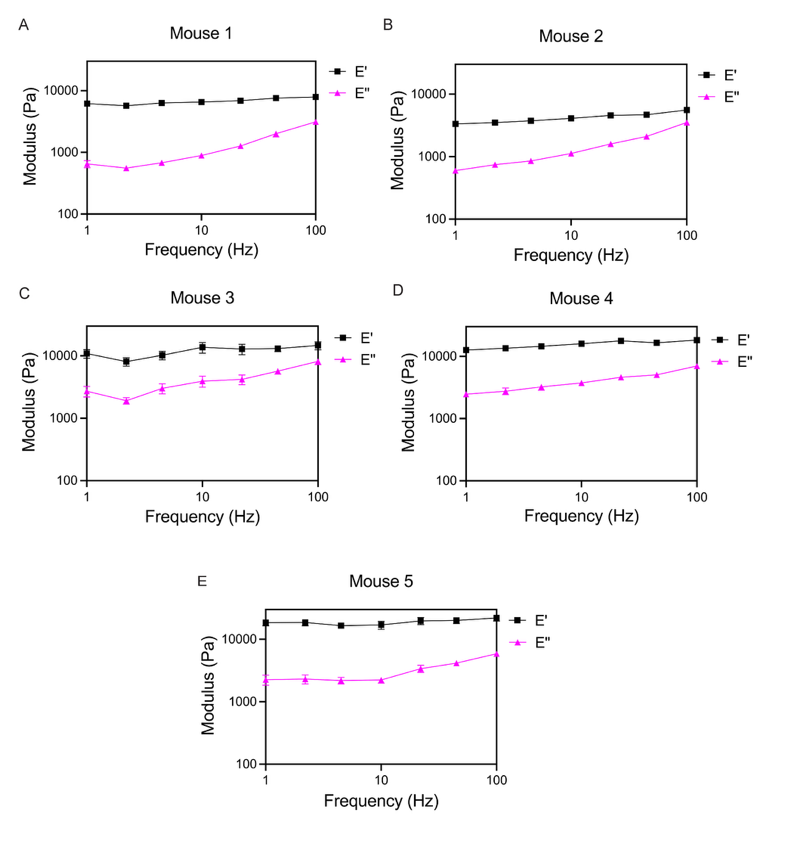

**Supplemental Figure 12: AFM frequency sweep of individual tumors.** (A-E) Frequency sweep for storage modulus E’ and loss modulus E’’ measured by AFM for PBS treated mice 1-5.

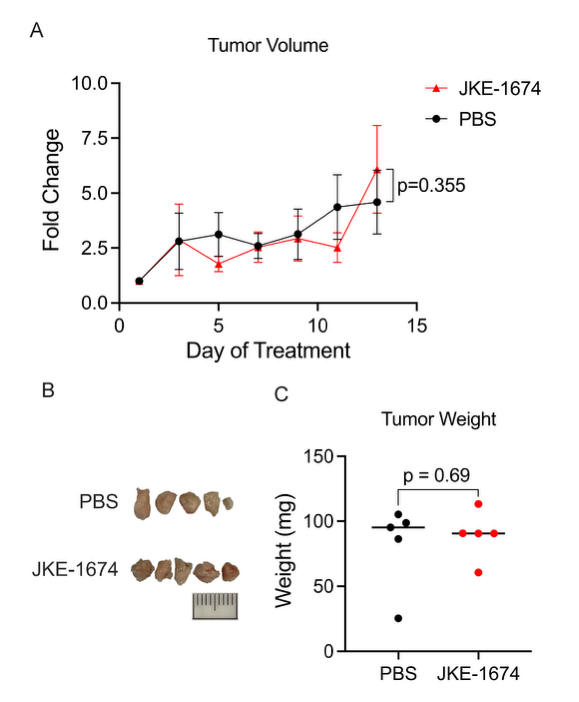

**Supplemental Figure 13: Ferroptosis lacks therapeutic potential for soft breast tumors.** (A) Fold change in tumor volume, measured by calipers, relative to the initial tumor volume on day 7 after treatment initiation. (B) Images of isolated tumors for each condition. Scale bar = 10 mm (C) Measured weight of whole isolated tumors for each condition. (N=5 for all conditions and measurements). Figure 1A graph represents means ± SEM. Lines in graph in C represent mean. One-way ANOVA was used for C. Two-way repeated-measures ANOVA was used for A.

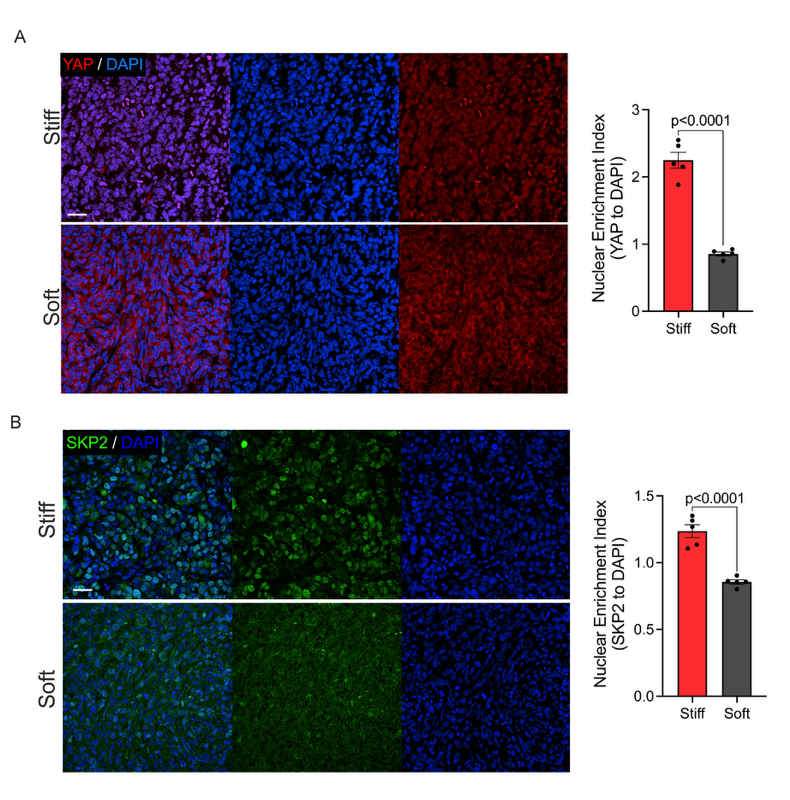

**Supplemental Figure 14: YAP and SKP2 stains for soft and stiffened tumors.** (A) Representative immunofluorescence images of MDA-MB-231 soft or stiff generated tumors stained with DAPI (nuclei in blue) and YAP (in red) (left) and quantification (right). Scale bar=25 $\mu m$. Representative image shown; analysis was performed on tumors from N = 5 mice per group. (B) Representative immunofluorescence images of MDA-MB-231 soft or stiff generated tumors stained with DAPI (nuclei in blue) and SKP2 (in green) (left) and quantification (right). Scale bar=25 $\mu m$. Representative image shown; analysis was performed on tumors from N = 5 mice per group. Nuclear enrichment index >1 indicates nuclear enrichment and <1 nuclear exclusion. All graphs represent means ± SEM. One-way ANOVA was used for analysis.

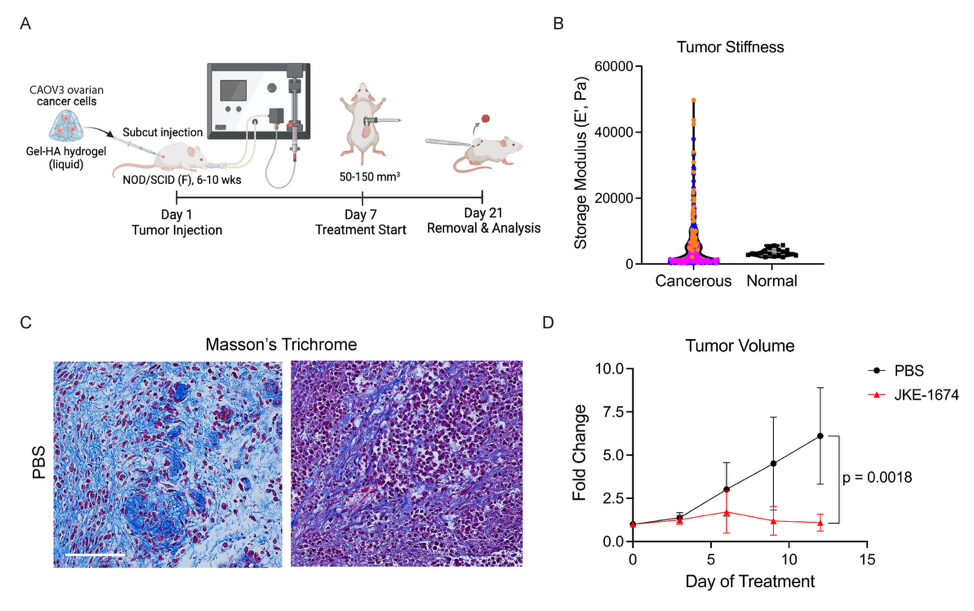

**Supplemental Figure 15: Ferroptosis can target stiff ovarian tumors.** (A) Schematic of experimental timeline for establishing stiff ovarian tumors in vivo. (B) AFM measurements of fresh ovarian tumor sections compared to normal ovarian tissue from (52). (C) Representative images of Masson’s trichrome staining of PBS treated tumor sections. Scale bars=100 $\mu m$ (D) Fold change in tumor volume, measured by calipers, relative to the initial tumor volume on day 7 after treatment initiation. (N=3) All graphs represent means ± SEM except B. Two-way repeated-measures ANOVA was used for D.
